# Temperature affects conspecific and heterospecific mating rates in *Drosophila*

**DOI:** 10.1101/2024.10.28.620639

**Authors:** Jonathan A. Rader, Daniel R. Matute

## Abstract

Behavioral mating choices and mating success are important factors in the development of reproductive isolation during speciation. Environmental conditions, especially temperature, can affect these key traits. Environmental conditions can vary across, and frequently delimit species’ geographic ranges. Pairing suboptimal conditions with relative rarity of conspecifics at range margins may set the stage for hybridization. Despite the importance of mating behaviors as a reproductive barrier, a general understanding of the interaction between behavioral choices and the environment is lacking, in part because systematic studies are rare. With this report, we begin to bridge that gap by providing evidence that temperature has a significant - but not consistent influence on mating choices and success, and thus on reproductive isolation in *Drosophila*. We studied mating propensity and success at four different temperatures among 14 *Drosophila* species in non-choice conspecific mating trials and in heterospecific trials among two *Drosophila* species triads that are known to regularly hybridize in the wild. We show that mating frequency varies significantly across a 10°C range (from 18ºC to 28ºC), both in 1:1 mating trials and in high-density *en-masse* trials, but that the effect of temperature is highly species-specific. We also show that mating frequency is consistently low and that temperature has a moderate effect in some heterospecific crosses. As conspecific mating propensity decreases outside of the optimal thermal range, while heterospecific matings remain constant, the proportion of heterospecific matings at suboptimal temperatures is relatively high. This result indicates that temperature can modulate behavioral choices that impose reproductive barriers and influence the rate of hybridization. More broadly, our results demonstrate that to truly understand how mating choice and reproductive isolation occur in nature, they need to be studied in an environmental context.

## INTRODUCTION

Speciation is the process in which one species splits into two lineages which in turn, accumulate genetic differences over time (Coyne and Orr, 2004). These lineages can persist through secondary contact when traits have diverged such that they preclude the formation of hybrids, or when accumulated genetic incompatibilities render hybrids less fit than the parental lineages. Thus, traits that hamper gene exchange between species are crucial for the maintenance of species boundaries. In animals, behavioral mating choices are an important component of speciation, particularly when they result in reproductive isolation (Kirkpatrick and Ravigné, 2002; Mendelson and Safran, 2021; Ritchie, 2007). Comparative studies have revealed that mating discrimination can emerge rapidly between species, and ensues faster than the evolution of postmating-prezygotic isolation (Turissini et al., 2018) and of postzygotic isolation (Coyne and Orr, 1997, 1989; Matute and Cooper, 2021; Turelli et al., 2014).

Behavioral isolation has therefore been hypothesized to evolve faster than any other trait involved in reproductive isolation among animals. Barriers to gene flow have often been described as intrinsic or extrinsic depending on whether they are affected by the environmental conditions in which species encounter each other (Coyne and Orr, 2004; Nosil, 2012). Nonetheless, all genetically-based traits, including those that are involved in reproductive isolation, also have an environmental component that interacts with the genome. This means that traits involved in reproductive isolation have interacting genetic and environmental components, and thus the gene × environment interactions might be a key aspect of species formation and persistence in the face of potential gene flow. Systematic surveys of the strength of different barriers to gene flow in different environments are rare but there is evidence that environmental conditions affect reproductive barriers. For example, immigrants can be less fit (or outright inviable) when they arrive in new environments (Giraud, 2006; Nosil et al., 2005). Additionally, environmental conditions are rarely homogenous across a species’ range. Conditions in the core of the range may be more suitable than at its periphery, and theory suggests that this, paired with the relative rarity of conspecifics, may lead to more frequent hybridization at the edge of the geographic range (Wilson and Hedrick, 1982). Similarly, postzygotic isolation barriers, developmentally-based defects that occur after a zygote is formed, have been conventionally considered intrinsic, but systematic studies have shown that environmental conditions can affect the magnitude and impact of these traits. For example, temperature determines the penetrance between negative epistatic interactions that reduce hybrid fitness between *Drosophila* species (Coyne et al., 1998; Miller and Matute, 2017; Presgraves, 2003). Similarly, hybrid newts experience greater oxidative stress at higher temperatures, a phenomenon not observed in the parental species (Petrović et al., 2023). Despite these examples of temperature influencing hybrid inviability, a general understanding about the magnitude of temperature effects in other barriers to gene flow is lacking. A particular aspect that remains understudied in behavioral isolation is how mating behaviors change along continuous environmental gradients, the most important of which might be temperature.

Environmental temperature has profound influences on most aspects of life in ectotherms, including numerous aspects of reproduction such as mating and reproductive success (Ingleby et al. 2010, Garcia-Roa et al. 2020, Leith et al. 2021). A recent meta-analysis examined how temperature changes influenced mating latency, choosiness and mating success, three of the components of mating choice in animals (Pilakouta and Baillet, 2022), and suggested an increase in mating success when animals are exposed to higher temperatures during mating trials but not when they were exposed before matings (Pilakouta and Baillet, 2022). This is surprising because for species that exist across an elevational or latitudinal gradient, the concomitant temperature gradient is frequently a key factor that delimits species’ geographic ranges. Assessments of the impact of temperature on behavioral isolation are thus important to assess the relative importance of different barriers to interbreeding (Leith et al. 2022).

To address this gap, we studied the effect that environmental conditions have on premating isolation in *Drosophila*. In particular, we studied mating rates in both conspecific and heterospecific pairings of 14 *Drosophila* species in different temperature regimes. We used non-choice experiments across a 10°C range (from 18°C to 28°C), observing the responses in both *en-masse* group experiments and 1:1 individual trials. We find that conspecific mate choice is highly contingent on the environmental temperature at which mating takes place, but heterospecific mating rate remains largely constant, and low. Two exceptions occurred among species that are known to hybridize (Comeault et al., 2016; Llopart et al., 2005; Matute and Ayroles, 2014; Schrider et al., 2018), with sexual isolation appearing to be less effective at low temperatures. These results suggest that environmental factors are important in modulating the strength of sexual isolation, with non-optimal environmental conditions perhaps fostering a higher risk of hybridization. Our results further suggest that behavioral choices are similar to other reproductive barriers in their contingency on environmental conditions.

## MATERIALS AND METHODS

### Species and Stocks

Our goal was to determine whether temperature affected mating rate in conspecific and heterospecific crosses of different species of *Drosophila*. We used fourteen *Drosophila* species. Three of the species were from the *yakuba* species complex (*D. yakuba*, *D. santomea*, *D. teissieri*), three from the *simulans* species complex (*D. sechellia*, *D*. *simulans, D. mauritiana*). We also used *D. melanogaster*, *D. pseudoobscura*, *D. willistoni*, *D. paulistorum*, *D. sturtevanti*, *D. virilis*, *D. americana*, and *D. novamexicana*. For all experiments, we used isofemale lines, shelf-stable stocks derived from the progeny from a single female collected in the field (David et al., 2005). Figure 1 shows the phylogenetic relatedness of the species in our sample (based on Kim et al, 2021 and Kim et al. 2024), and which species show range overlap in their natural ranges (based on Yukilevich 2012). Table S1 shows the origin, collector, and other details of each isofemale line used in this study. All isofemale lines were maintained in triplicate on corn-meal substrate in 30mL plastic vials since their establishment. Prior to experiments, parallel stocks of flies were reared and maintained in incubators at each of four temperatures (18°C, 22°C, 25°C, and 28°C). We only used one isofemale line per species, and while this limits our ability to assess intraspecific variation, the scale of the experiments did not allow us to include more isofemale lines and remains sufficient to study interspecific effects - the focus of the present work. While previous assessments have detected variation in mating behavior in some of the included species (e.g., Coughlan et al. 2022, Li et al. 2023, Matute et al. 2010), our goal was to assess the existence of variation in mating behavior, and not quantify the magnitude of the effect of temperature in mating behavior in different genetic backgrounds. The latter goal would require a much larger experiment.

**FIGURE 1.**
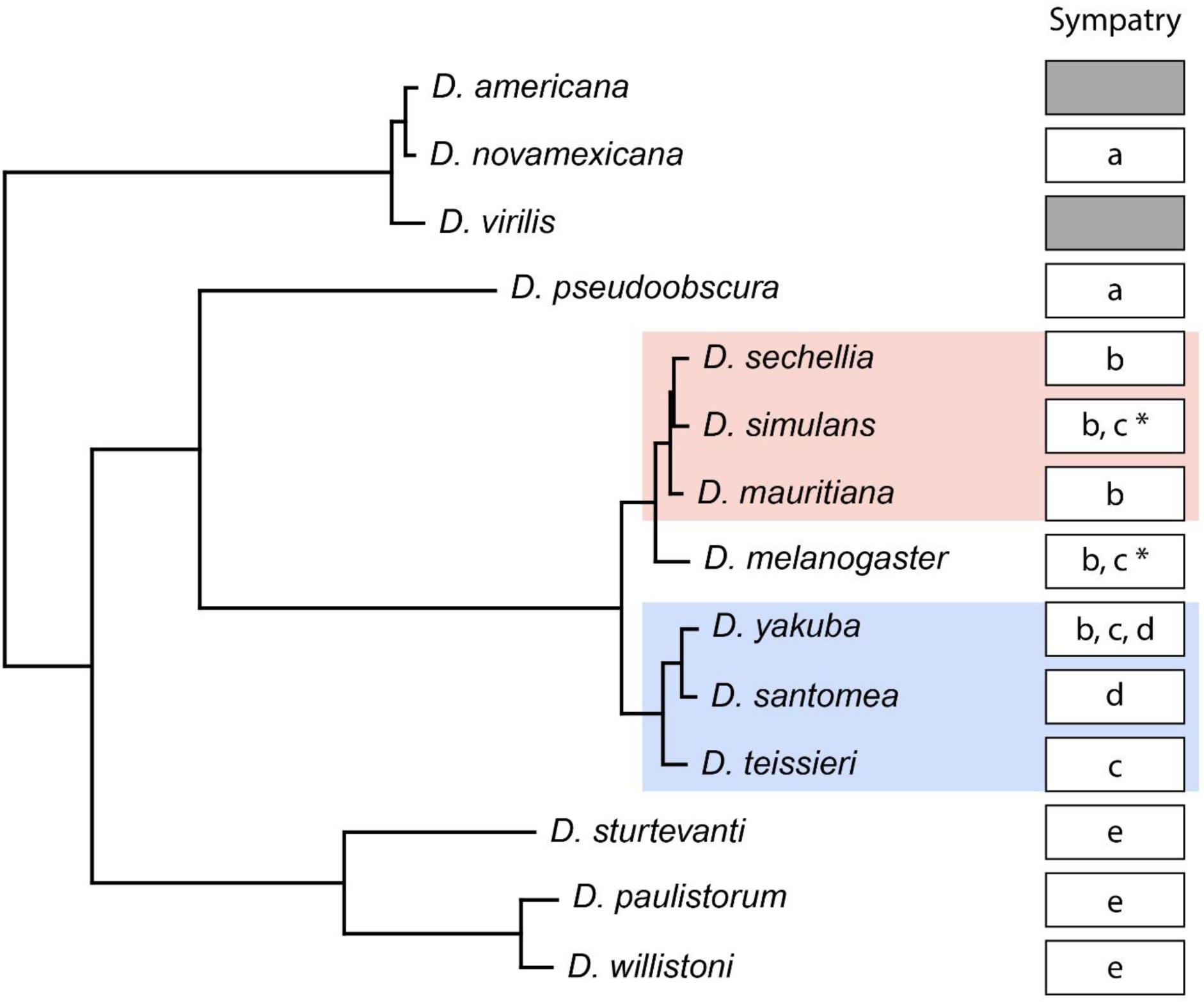
Phylogeny and sympatry of Drosophila species included in this study. The phylogeny here was pruned from the Kim et al. (2021, 2024) phylogeny to include the 14 species in our mating choice trials. These include two recently-diverged species triads with species that are known to hybridize in the wild (red and blue shaded regions). Boxes show which species share overlapping geographic ranges.

Among these 14 species, several species pairs show the ability to produce hybrids. The three species from the *virilis* phylad (*D. virilis*, *D. americana*, and *D. novamexicana*) produce fertile offspring in all the reciprocal directions (Hoikkala and Lumme 1987; reviewed in Yukilevich 2012). Genome assessments have also revealed the existence of introgression among these species (Yusuf et al. 2022) but to date, no hybrid zone has been identified for this group. The same is true for the species in the *willistoni* species group (*D. willistoni* and *D. paulistorum*) which show evidence of hybridization (Carson 1954, Dobzhansky and Spassky 1959, Winge and Cordeiro 1965), and introgression (Suvorov et al. 2022) but no known stable hybrid zones. Species from the *melanogaster* species group also show evidence of hybridization (reviewed in Turissini et al. 2018) and introgression (Turissini and Matute 2017, Schrider et al. 2018, Suvorov et al. 2022). Three species pairs form stable hybrid zones in the wild: *D. santomea* with *D. yakuba* (Llopart et al. 2004, Matute et al. 2010, Comeault et al. 2016), *D. simulans* with *D. sechellia* (Matute and Ayroles 2014, Schrider et al. 2018) and *D. yakuba* with *D. teissieri* (Cooper et al. 2018). Hybridization is not known from the other species in our sample, despite widespread sympatry (see Figure 1). To understand the effects of temperature on interspecific mating behavior, we focused on the melanogaster species group, given their occurrence of hybrid zones in nature. We studied the two reciprocal crosses of six hybridizations: *D. santomea* × *D. yakuba*, *D. teissieri* × *D. yakuba*, *D. santomea* × *D. teissieri*, *D. simulans* × *D. sechellia*, *D. simulans* × *D. mauritiana*, and *D. sechellia* × *D. mauritiana*.

### Virgin Collection

About one month before starting experiments, we expanded each isofemale line for the study from 3mL vials into 100mL plastic bottles with cornmeal and yeast (35 g/L, Red Star active dry yeast, *Saccharomyces cerevisiae*). As soon as we observed larvae in the food, we removed the adults, added 1 ml of propionic acid (0.5% v/v) solution to the bottles, and provided a pupation substrate (Kimwipes Delicate Task; Kimberly Clark, Irving, TX). Once we observed black pupae in the KimWipes or the walls of the bottle, we cleared the bottle by transferring all adults to a different bottle. Then, we inspected the bottles every 8 hours and collected any adults that had hatched in that period of time. To separate virgins, we anesthetized all the flies in each bottle on a Flypad with CO_2_. Flies were then placed in groups of 20 individuals (separated by sex) in 30mL cornmeal vials.

### Mating Experiments

We ran two different types of mating experiments: (*i*) non-choice *en-masse* experiments, and (*ii*) individual non-choice experiments. We describe the rationale and the procedure for each of them as follows.

#### (i) Non-choice *en-masse*

First, we studied the frequency of conspecific matings (i.e., females and males from the same species) in *en-masse* experiments. Briefly, we collected females as described immediately above. We combined 100 females and 100 males from either conspecific or heterospecific lines in a 30mL vial and housed each female-male combination for 24 hours at each of the 4 temperatures in which they were reared: 18°C, 22°C, 25°C, and 28°C, respectively. For each conspecific, we did five replicates for a total of 280 conspecific *en-masse* matings (14 species × 4 temperature × 5 replicates). After 24h, females were separated using CO_2_ anesthesia and the reproductive tract of each female was removed using tweezers. Batches of 5 to 10 reproductive tracts were mounted in Ringer’s solution (NaCl, KCl, CaCl_2_ and NaHCO_3_ in distilled water). We inspected both the spermatheca and the seminal receptacle to assess the presence of sperm, which is indicative of mating (Turissini et al., 2018). The proportion of mated females per vial was a proxy of the extent of sexual isolation.

To determine whether temperature affected the likelihood of mating in conspecific crosses, we used two different regression families. First, we fit a binomial logistic regression in which female mating status was the response, species was a fixed effect, and temperature was a continuous effect. The model also included the interaction between sex and temperature. We used the R function *glm* (family = “binomial” (library *stats*, R Core Team, 2018) to fit the models. To determine the significance of each effect, we used likelihood ratio tests using the function *lrtest (library lmtest,* Hothorn et al., 2015) in which we compared a model with and without the effect to be tested. The logistic linear model followed the form:

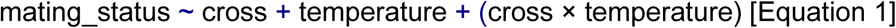

We also fit a logistic regression that included a quadratic temperature term and the interaction with cross. The model followed the form:

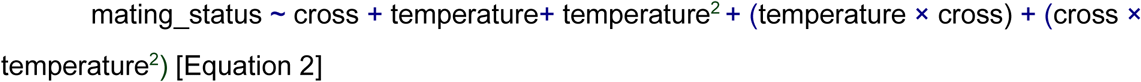

Second, we studied the frequency of heterospecific matings (i.e., females and males from different species), also in an *en-masse* setting. Table S2 lists the possible interspecific crosses included in this portion of the research. For each heterospecific mating, we did five replicates for a total of 60 interspecific assays (6 crosses × 2 directions × 5 replicates). We scored the proportion of mated females in the same way as described for conspecific matings (immediately above).

For each type of interspecific cross, we fit a binomial regression in which the mating status of the dissected female was the response, the direction of the cross was a fixed effect, and temperature was a continuous effect. The model also included the direction × temperature interaction. The linear logistic model had the following form:

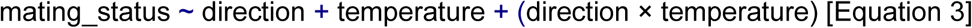

We also fitted a second model including the quadratic temperature term which took the form:

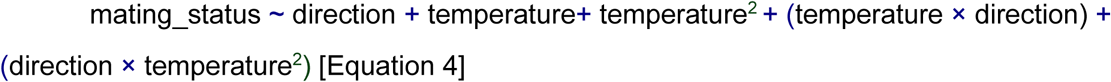

In total, we fit twelve regression models for these analyses (six linear and six quadratic). We compared the model fit using AIC values (calculated using the function AIC, library stats, R Core Team 2018) and only retain the quadratic model if AIC_Quadratic_ had a better value than the AIC_Linear_ by two AIC units. Every model described here (conspecific or heterospecific), was followed by posthoc Tukey HSD tests using the function *glht* (library *multcomp*, (Hothorn et al., 2016, 2008)).

#### (ii) Non-choice individual pairs

We also conducted experiments in which we watched pairs of flies (one male and one female) to study two key characteristics of sexual behavior, namely copulation latency (the time that it takes for copulation to begin) and copulation duration (the time that coupling lasts). We measured copulation latency and duration in both conspecific and heterospecific crosses using non-choice mating experiments. Matings all were run in a climate-controlled room at four different temperatures (18°C, 22°C, 25°C, and 28°C) corresponding with the temperatures at which the flies were reared. All flies in this experiment were collected as virgins and housed in single sex vials as described in the section immediately above (Virgin collection). On day four after hatching, one female and one male were aspirated into a single, empty vial. All mating trials were started within 1 hr of the beginning of the light cycle to maximize fly activity and female receptivity. We observed 100 pairs per genotype pairing, and the flies were watched constantly for 1 hour. No more than 200 vials were set up in parallel to ensure accuracy in recording when copulation began and ended. For each of the pairs, we recorded whether mating took place, and in cases where copulation occurred, we timed latency (the time to copulation initiation) and duration (time from mounting to separation).

To determine whether temperature had an effect on mating frequency in conspecific matings, we fit a linear and quadratic logistic models identical to the ones we used for *en-masse* experiments (Equations 1 and 2). We also use AIC values to determine which model was a better fit to the data.

We compared the results of the individual trials to those of the *en-masse* experiments using a set of Pearson’s correlation tests (function *corr*, library *stats*, R Core Team 2018) of mating rate in *en-masse* trials vs. mating rate in individual trials at each of the four test temperatures. We also fit quadratic models to study the effect of temperature in conspecific mating latency and duration. The models were identical to those presented in equations 1 and 2, except that they were not logistic but gaussian models.

We followed a similar approach to analyze data from heterospecific matings. For each hybridization in which we obtained latency and duration data from both directions of the cross (five hybridizations), we fit a linear model for each of the mating traits with the direction of the cross as a fixed effect, and temperature as a continuous effect, and included an interaction between these two effects (i.e., a model similar to equation 3 but with a gaussian distribution instead of a logistic one). Similarly, we conducted model fit assessments using the models’ AIC values. The cross *D. sechellia* × *D. mauritiana* yielded no matings during the 1-hour observation period at any of the temperatures, thus we did not conduct statistical analyses for this hybridization.

Finally, we used the rates of mating in conspecific matings and heterospecific matings to calculate an index of sexual isolation for each direction of interspecific crosses. The index, *I_S_*, normalizes the proportion of heterospecific matings in a direction by the number of matings in conspecific crosses involving the same female genotype.

The index follows the form:

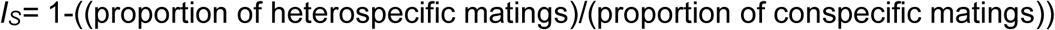

*I_s_* usually ranges between 0 (when heterospecific matings are as common as conspecific matings) and 1 (when heterospecific matings do not occur), however negative values can occur and indicate that heterospecific matings are more common than the conspecifics. We used the Agresti-Coull Interval Method to calculate the confidence intervals for the heterospecific and conspecific mating proportions (function *add4ci*, library *PropCIs*, Scherer and Scherer, 2018) and a 2-sample test for equality of proportions with continuity correction (function *prop.test*, library *stats*, R Core Team, 2018) to compare mating proportions.

### Ethical Note

The species in this study do not require licenses or permits. Mated females were dissected under CO_2_ anesthesia to minimize pain. All flies were killed by immersion in isopropanol after experimentation.

## RESULTS

### Temperature effect in conspecific and heterospecific *en-masse* matings

First, we studied whether environmental temperatures affected the likelihood of mating between conspecifics in *en-masse* matings. Figure 2A shows the relative frequency of mating for 14 species at four temperatures. A logistic regression with binomial responses on the *en-masse* experiments conducted at different temperatures revealed that environmental temperature had a strong effect on whether mating took place, but also strong heterogeneity among species suggests differential effects of temperature across species. Notably, we find a strong interaction between temperature and species identity suggesting that the effect of temperature varies among species and that its magnitude is not universal (Table 1). Adding a quadratic temperature effect also showed strong effects of species, temperature, and the interaction between these two effects, but also revealed a significant contribution of the squared temperature, and the interaction between the squared temperature and species (Table 1). Notably, the addition of the quadratic term improves the model fit (AIC_Linear_ = 36,412.97; AIC_Quadratic_= 33,057.5), indicating that the likelihood of conspecific mating is better explained by a nonlinear model. Figure S1 shows the proportion of mated females for each species along the temperature continuum.

**FIGURE 2.**
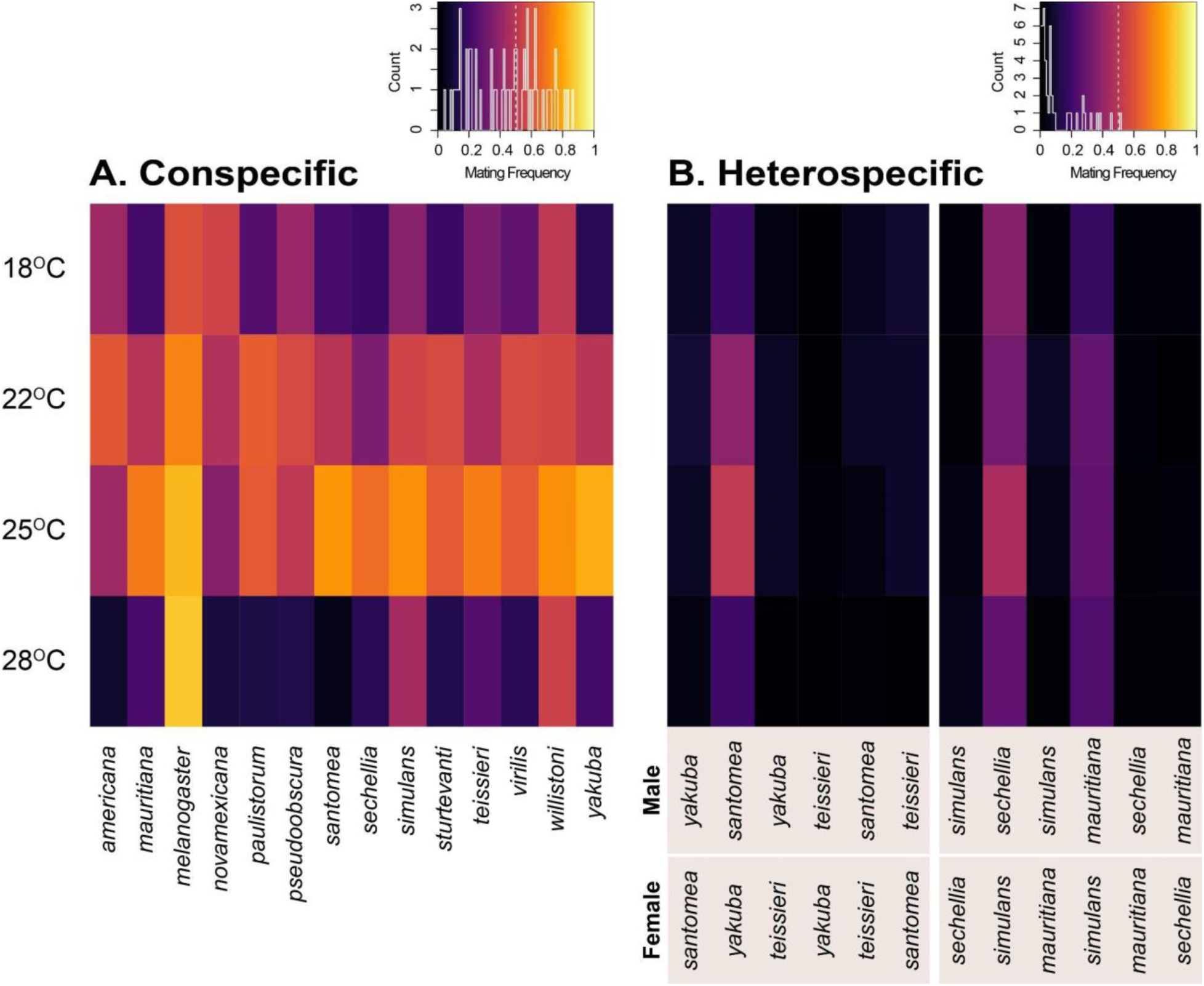
Mating frequency in *en-masse* mating experiments. **A.** Mating frequency in conspecific pairings of 14 different species of *Drosophila*, measured by counting inseminated females from groups of 100 females and 100 males. **B.** Mating frequency in reciprocal heterospecific matings between six *Drosophila* species pairs, similarly measured as inseminated females.

**TABLE 1.** Temperature affects the likelihood of conspecific mating in enmasse settings. We show the model fit for the linear and quadratic models.

|  | Linear |  |  | Quadratic |  |  |
| --- | --- | --- | --- | --- | --- | --- |
|  | LRT | df | *P* | LRT | df | *P* |
| Species | 13.472 | 1 | $2.421 \times 10^{-4}$ | 1,467.21 | 13 | $< 1 \times 10^{-10}$ |
| Temperature | 1,320.749 | 1 | $< 1 \times 10^{-10}$ | 2,828.02 | 1 | $< 1 \times 10^{-10}$ |
| Temperature<sup>2</sup> | NA | NA | NA | 2,859.67 | 1 | $< 1 \times 10^{-10}$ |
| Species x temperature | 613.909 | 13 | $< 1 \times 10^{-10}$ | 521.82 | 13 | $< 1 \times 10^{-10}$ |
| Species × temperature<sup>2</sup> | NA | NA | NA | 523.80 | 13 | $< 1 \times 10^{-10}$ |

Next, we studied whether temperature had an effect on the frequency of heterospecific matings in the *en-masse* setting. For these analyses, we also used linear and quadratic and logistic regressions. In five of the hybridizations, the quadratic model fit better than the linear model (AIC values listed in Table S3). The only exception was the *D. mauritiana* × *D. sechellia* crosses in which the linear model was a better fit. In four of the hybridizations (*D. yakuba* × *D. santomea*, *D. yakuba* × *D. teissieri*, *D. santomea* × *D. teissieri*, and *D. simulans* × *D. mauritiana*), either the temperature effect or the square temperature effect were significant, indicating that the likelihood of heterospecific mating in two directions of the cross was affected by temperature (Table 2 and Table S4). The same four hybridizations showed a significant interaction between the square of the temperature and the direction of the cross. Two crosses also showed a significant interaction between temperature and cross direction. The significance of these interactions indicates a differential effect of temperature in the two directions of the cross in these hybridizations. Notably, these interactions are significant in the three hybridizations of the *yakuba* species group, a group of species with marked interspecific differences in thermal fitness (Comeault and Matute, 2021; Cooper et al., 2018; Matute et al., 2009). Table S4 shows the results for the linear models, which also show a species-dependent effect of temperature, but suggest a much stronger effect of the reciprocal cross direction. The crosses *D. mauritiana* × *D. sechellia* showed no strong effect of temperature in the linear or quadratic analyses, but do note that the number of matings in these two reciprocal crosses was low.

**TABLE 2.** Quadratic logistics regressions suggest a common effect of temperature in the likelihood of heterospecific mating in *en-masse* experiments. The metric of isolation is receptivity of females in *en-masse* matings. All the likelihood ratio tests (LRT) comparisons involve one degree of freedom.

| Species pair | Direction |  | Temperature |  | Direction x temperature |  | Temperature<sup>2</sup> |  | Temperature<sup>2</sup> x direction |  |
| --- | --- | --- | --- | --- | --- | --- | --- | --- | --- | --- |
| | $\chi^2$ | *P* | $\chi^2$ | *P* | $\chi^2$ | *P* | $\chi^2$ | *P* | $\chi^2$ | *P* |
| *D. yakuba*/<br>*D. santomea* | 7.238 | 0.007 | **10.527** | **1.177 × 10<sup>-3</sup>** | **7.836** | **5.122 × 10<sup>-3</sup>** | **11.532** | **6.841 × 10<sup>-4</sup>** | **6.360** | **0.012** |
| *D. yakuba*/<br>*D. teissieri* | 0.003 | 0.958 | **24.299** | **8.249 × 10<sup>-7</sup>** | 0.046 | 0.830 | **25.419** | **4.613 × 10<sup>-7</sup>** | <u>0.067</u> | <u>0.796</u> |
| *D. santomea*/<br>*D. teissieri* | 0.003 | 0.955 | **12.267** | **4.612 × 10<sup>-4</sup>** | 0.014 | 0.905 | **14.509** | **1.395 × 10<sup>-4</sup>** | <u>0.014</u> | <u>0.907</u> |
| *D. simulans*/<br>*D. sechellia* | 1.682 | 0.195 | 1.342 | 0.247 | 3.739 | 0.0532 | 1.764 | <u>0.184</u> | **4.564** | **0.0327** |
| *D. simulans*/<br>*D. mauritiana* | 7.060 | 0.008 | **18.214** | **1.975 × 10<sup>-5</sup>** | **6.091** | **0.014** | **18.799** | **1.452 × 10<sup>-5</sup>** | **6.763** | **9.309 × 10<sup>-3</sup>** |
| *D. sechellia*/<br>*D. mauritiana* | 3.190 × 10<sup>-4</sup> | 0.986 | 0.033 | 0.855 | 1.058 × 10<sup>-3</sup> | 0.974 | 0.095 | <u>0.755</u> | **7.973 × 10<sup>-3</sup>** | <u>0.929</u> |

### Temperature effect in individual non-choice matings

*En-masse* mating experiments allow rapid quantification of the rate of insemination in a combination of genotypes but make the measurement of individual mating components a challenge. We thus set up individual non-choice experiments (i.e., one male and one female). First, we studied whether temperature had an effect on mating success, copulation latency, and copulation duration in individual conspecific non-choice experiments. Figure 3 shows the mating frequency, latency, and duration in conspecific matings across 14 *Drosophila* species. Similar to our results in the *en-masse* experiments, both linear and quadratic models revealed that the likelihood of mating is affected by temperature, species identity, and the interaction between temperature and species. For mating frequency, the quadratic model shows a better fit than the linear model (AIC_Quadratic_ = 6,555.599, AIC_Linear_ = 7,136.356), and revealed a non-linear effect of temperature and a strong interaction between the quadratic term and species identity (Table 3). These results are similar to those of the *en-masse* experiments, which is not surprising given that the mating frequency in these individual non-choice experiments was highly correlated with the mating frequency in the non-choice *en-masse* experiments described immediately above (Figure 4, Table S5). Table S6 shows linear and quadratic regressions for the mating frequency of each species independently, which indicates that temperature was a significant effect in the best fitting model for 12 of 14 species.

**FIGURE 3.**
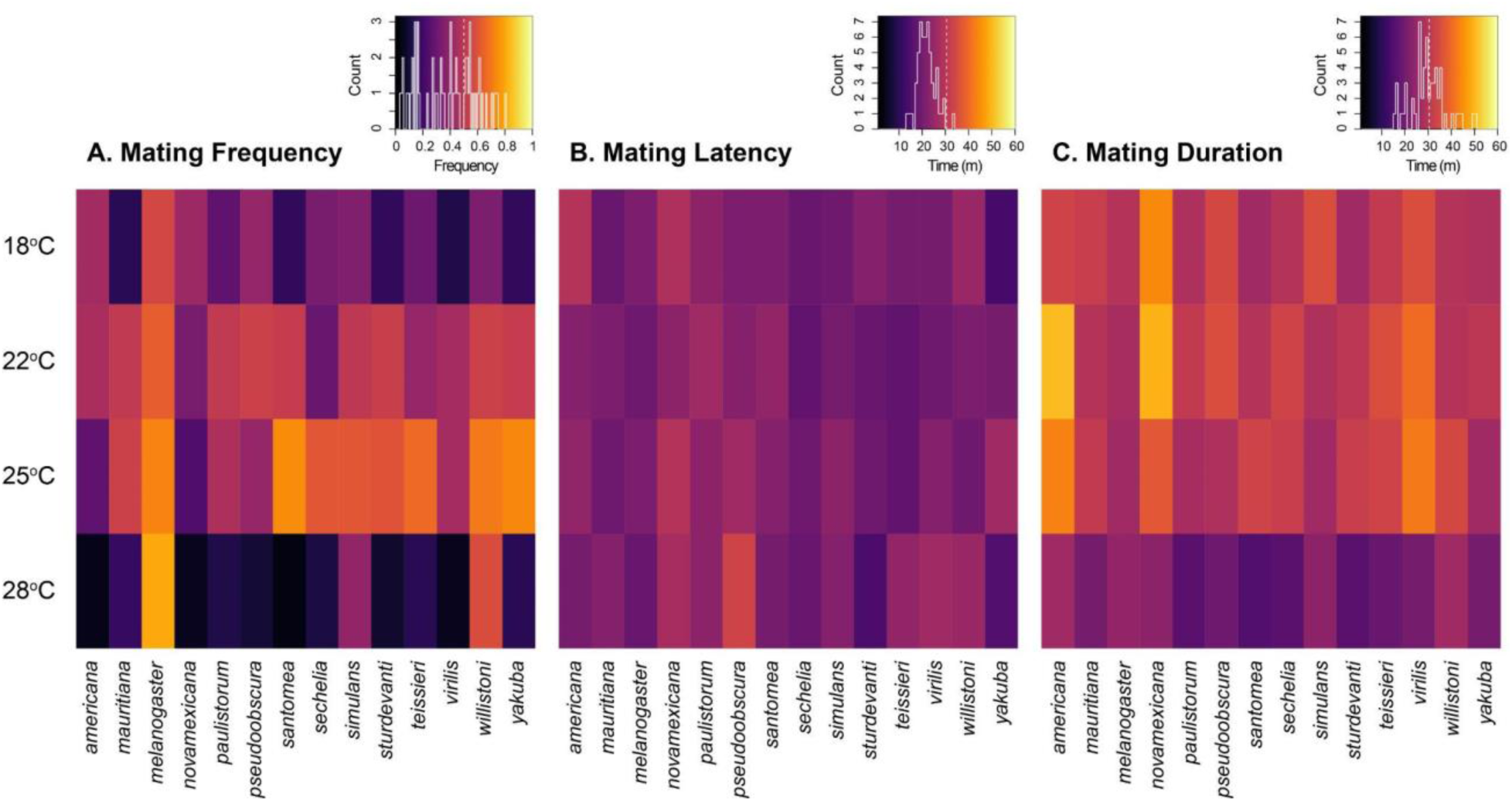
Non-choice individual experiments show strong differences in conspecific mating propensity at different temperatures. **A.** Mating frequency in conspecific mating trials of 14 different species of *Drosophila* measured as a proportion of successful matings over total trials. **B.** Mating latency in the same mating trials, measured as the time (in minutes) from the beginning of the trial to the onset of copulation. **C.** Mating duration in the individual trials, measured as the time (minutes) from mounting to separation.

**TABLE 3.**
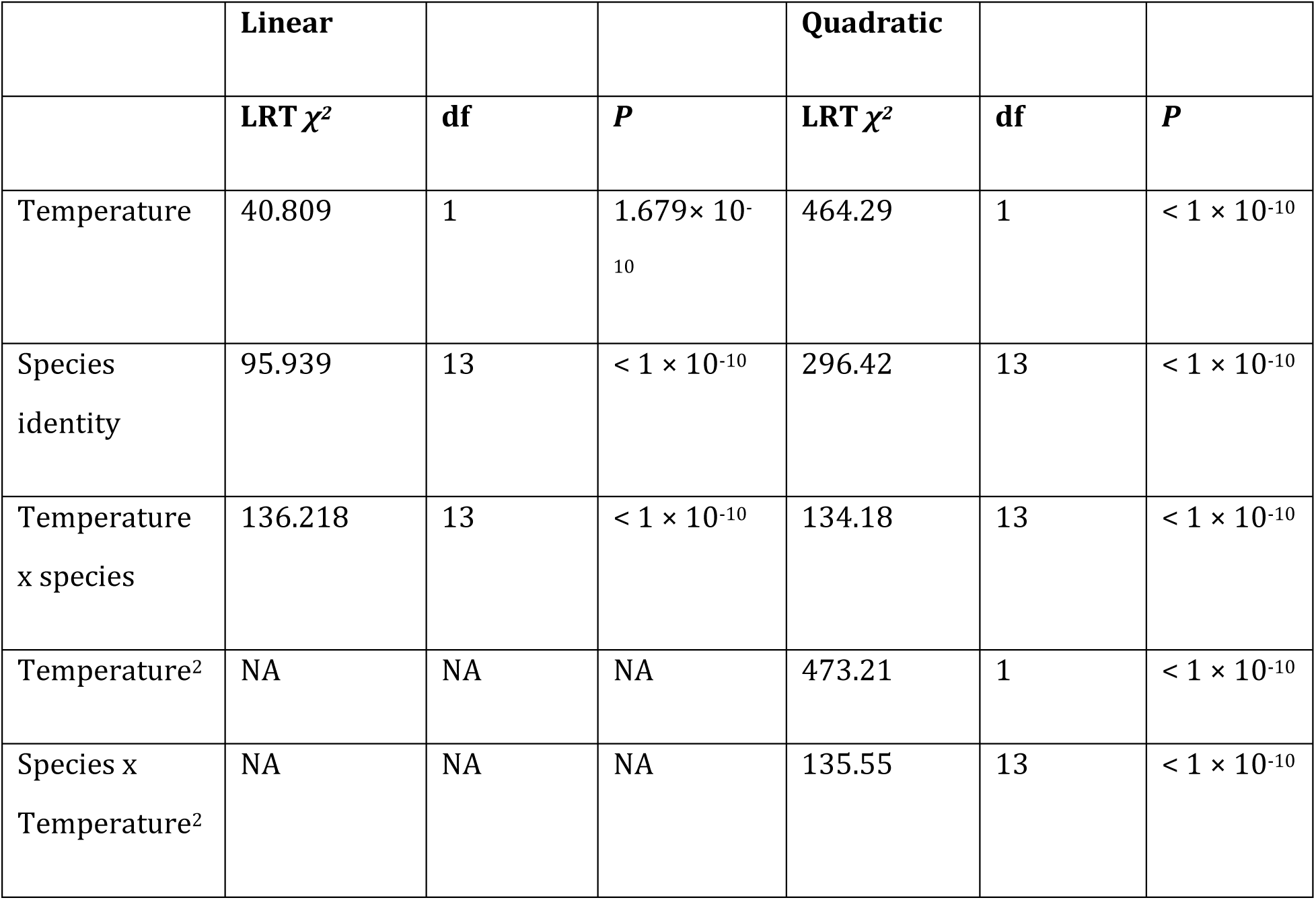
Temperature affects the likelihood of conspecific mating in individual settings. We show the model fit for the linear and quadratic models. The significance of each effect was determine with Likelihood ratio tests.

**FIGURE 4.**
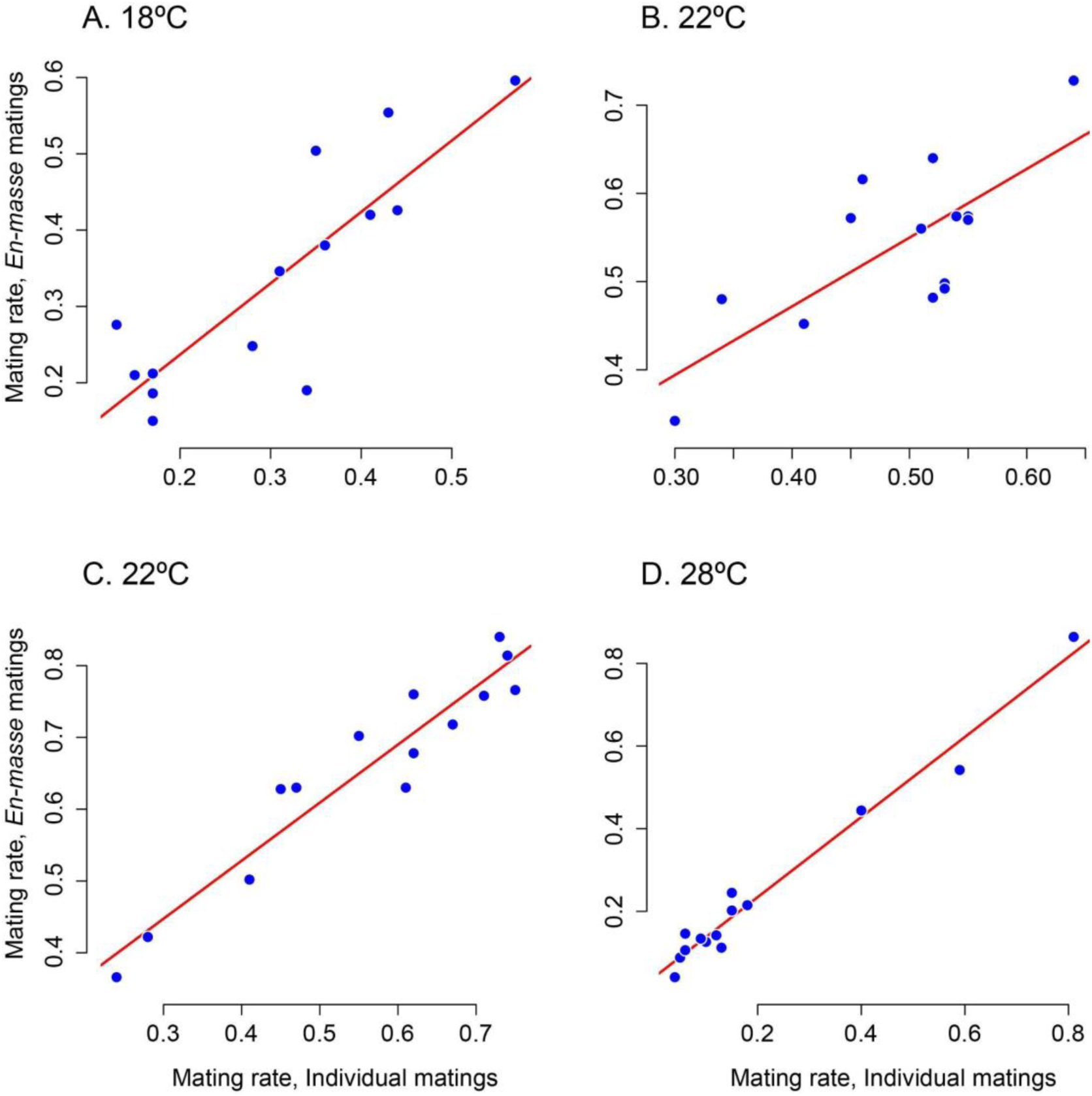
Mating rates are highly correlated between *en-masse* and individual non-choice experiments in conspecific matings. The four panels show correlations at four different temperatures. **A.** 18°C. **B.** 28°C. **C.** 25°C. **D.** 28°C.

To quantify the individual components of conspecific mating behavior, we fit linear and quadratic models to identify sources of heterogeneity in mating latency and mating duration in non-choice individual conspecific crosses. The addition of a quadratic term did not improve the model fit for conspecific copulation latency (AIC_Quadratic_ = 17,089.44, AIC_Linear_ = 17,088.44). Mating latency, or the time that it takes for mating to start, did not differ across species (Species effect: *F*_13,2163_ = 1.2972, *P* = 0.2067), and was not influenced by temperature (Temperature effect: *F*_1,2163_ = 2.2835, *P* = 0.1309, Figure 3B). There was no species-specific effect of temperature either (temperature × sex interaction: *F*_1,2163_ = 1.403, *P* = 0.150). These results indicate no generalized across-species effect of temperature on conspecific mating latency. Table S7 shows linear and quadratic regressions for the mating latency of each conspecific cross independently which indicates that temperature has some effect in three species, *D. yakuba*, *D. teissieri, D. melanogaster*, and *D. willistoni*. Figures S2 and S3 shows the mating latency for each species along the temperature continuum.

Mating duration in conspecific crosses, unlike latency, was better explained by a quadratic model than by a linear one (AIC_Quadratic_ = 17,979.11; AIC_Linear_ = 18,045.79). Duration was affected by species identity, temperature, the quadratic form of temperature, and the two interactions between cross and temperature (Table 4). These results suggest that while temperature had no detectable effect on mating latency in conspecific crosses, mating duration was influenced by temperature in a species-specific manner. Table S8 shows the fit for linear and quadratic regressions for each of the 14 species. Temperature influenced mating duration in all the best-fitting regressions (Table S8), suggesting that conspecific mating duration was affected by temperature in all species. Figures S4 and S5 shows the mating duration for each species along the temperature continuum.

**TABLE 4.** The identity of the cross and temperature have an effect on copulation duration of conspecific crosses.

| Effect | df | *F* | *P* |
| --- | --- | --- | --- |
| temperature<sup>2</sup> | 1 | 24.410 | $8.386 \times 10^{-7}$ |
| temperature | 1 | 26.315 | $3.161 \times 10^{-7}$ |
| cross | 13 | 3.274 | $5.950 \times 10^{-5}$ |
| temperature<sup>2</sup> x cross | 13 | 3.066 | 0.0001606 |
| Temperature x cross | 13 | 3.241 | $6.965 \times 10^{-5}$ |

Next, we studied the effect of temperature on the likelihood of mating in non-choice individual heterospecific assays. The quadratic models showed a better fit than linear models in two species pairs (*D. yakuba* × *D. santomea* and *D. simulans* × *D. mauritiana*, Table S9). Neither linear logistic nor quadratic logistic models show a generalized effect of temperature (Table 5 and Table S10). Unlike our analyses using *en-masse* matings, we found no significant interaction between cross direction and temperature in any of the hybridizations (Table 5) indicating differences in results depending on the experimental approach to study mating behavior in heterospecific crosses. Note that these experiments had fewer observations than *en-masse* experiments and that we observed no copulations for the ♀*D. sechellia* × ♂*D. mauritiana* cross at any temperature in this assay.

**TABLE 5.**
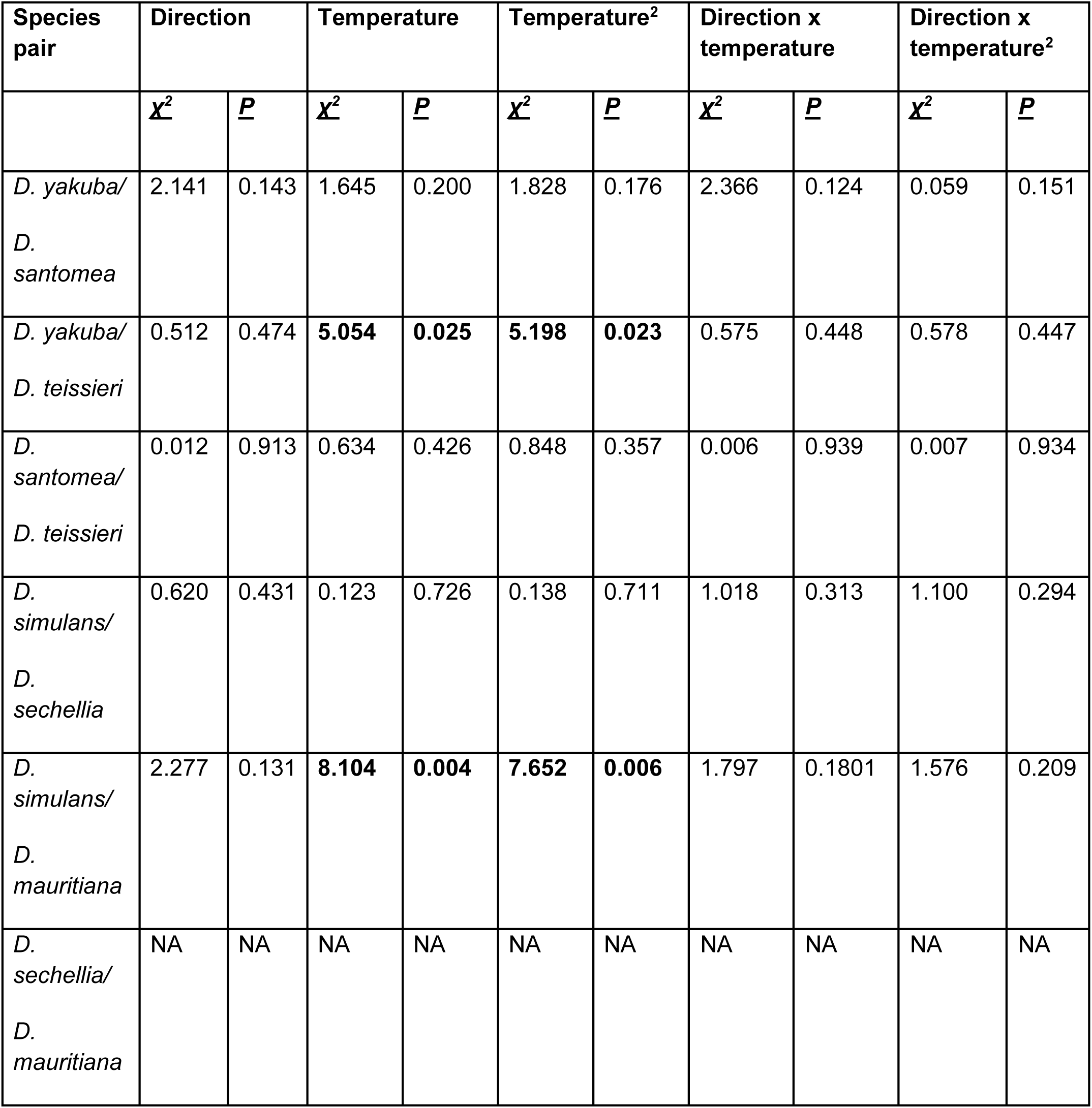
Temperature has a moderate effect on mating propensity in some of the six types of heterospecific *individual* matings. The metric of isolation is receptivity of females in *en-masse* matings. All the likelihood ratio tests (LRT) comparisons involve one degree of freedom. Quadratic logistics.

| Species pair | Direction |  | Temperature |  | Temperature<sup>2</sup> |  | Direction x temperature |  | Direction x temperature<sup>2</sup> |  |
| --- | --- | --- | --- | --- | --- | --- | --- | --- | --- | --- |
| | $\chi^2$ | *P* | $\chi^2$ | *P* | $\chi^2$ | *P* | $\chi^2$ | *P* | $\chi^2$ | *P* |
| *D. yakuba*/<br>*D. santomea* | 2.141 | 0.143 | 1.645 | 0.200 | 1.828 | 0.176 | 2.366 | 0.124 | 0.059 | 0.151 |
| *D. yakuba*/<br>*D. teissieri* | 0.512 | 0.474 | **5.054** | **0.025** | **5.198** | **0.023** | 0.575 | 0.448 | 0.578 | 0.447 |
| *D. santomea*/<br>*D. teissieri* | 0.012 | 0.913 | 0.634 | 0.426 | 0.848 | 0.357 | 0.006 | 0.939 | 0.007 | 0.934 |
| *D. simulans*/<br>*D. sechellia* | 0.620 | 0.431 | 0.123 | 0.726 | 0.138 | 0.711 | 1.018 | 0.313 | 1.100 | 0.294 |
| *D. simulans*/<br>*D. mauritiana* | 2.277 | 0.131 | **8.104** | **0.004** | **7.652** | **0.006** | 1.797 | 0.1801 | 1.576 | 0.209 |
| *D. sechellia*/<br>*D. mauritiana* | NA | NA | NA | NA | NA | NA | NA | NA | NA | NA |

From these individual heterospecific non-choice experiments, we obtained latency and duration measurements for eleven of the twelve interspecific crosses, or five bidirectional hybridizations and one unidirectional cross. In the five cases for which we had data for the two reciprocal crosses, the inclusion of quadratic terms did not improve the models fit to the latency or duration data. Tables S11 and S12 show the AIC values for the linear and quadratic models for heterospecific mating latency and duration, respectively. Linear models fit better for both latency and duration in all hybridizations. Mating latency in crosses between *D. simulans* and *D. sechellia* was the only instance in which the direction of the cross, temperature and the interaction between these two effects were significant (Table 6). These results indicate that temperature had no across-species effect on mating latency or duration in heterospecific crosses in *Drosophila* in timed individual mating trials.

Finally, we calculated an index of sexual isolation (*I_S_*) as a proxy of the strength of reproductive isolation resulting from mating choice at each temperature. This index represents the proportional risk of hybridization when normalizing by the number of conspecific matings that occur at any given temperature. Figure 6 shows the effect of temperature on isolation for *en-masse* and individual mating experiments. Sexual isolation is complete, or almost complete, in some crosses (i.e., ♀*D. sechellia* × ♂*D. mauritiana* and ♀*D. yakuba* × ♂*D. teissieri*), regardless of the temperature. On the other hand, the crosses between ♀*D. yakuba* × ♂*D. santomea* and between ♀*D. simulans* × ♂*D. sechellia* (and to a lesser extent ♀*D. santomea* × ♂*D. yakuba*) showed stark differences in total sexual isolation across temperatures with heterospecific matings being as common as conspecific at 18 and 28°C, the extreme values of the assayed temperature range. Despite finding some mean *I_S_* values < 0, we did not find any significant instances where the proportion of heterospecific matings exceeded the proportion of conspecific matings (*χ^2^* < 2.602, df = 1, *P* > 0.1067). These results indicate that the magnitude of sexual isolation as a barrier to gene flow might be contingent upon temperature, at least in some interspecific crosses.

**FIGURE 5.**
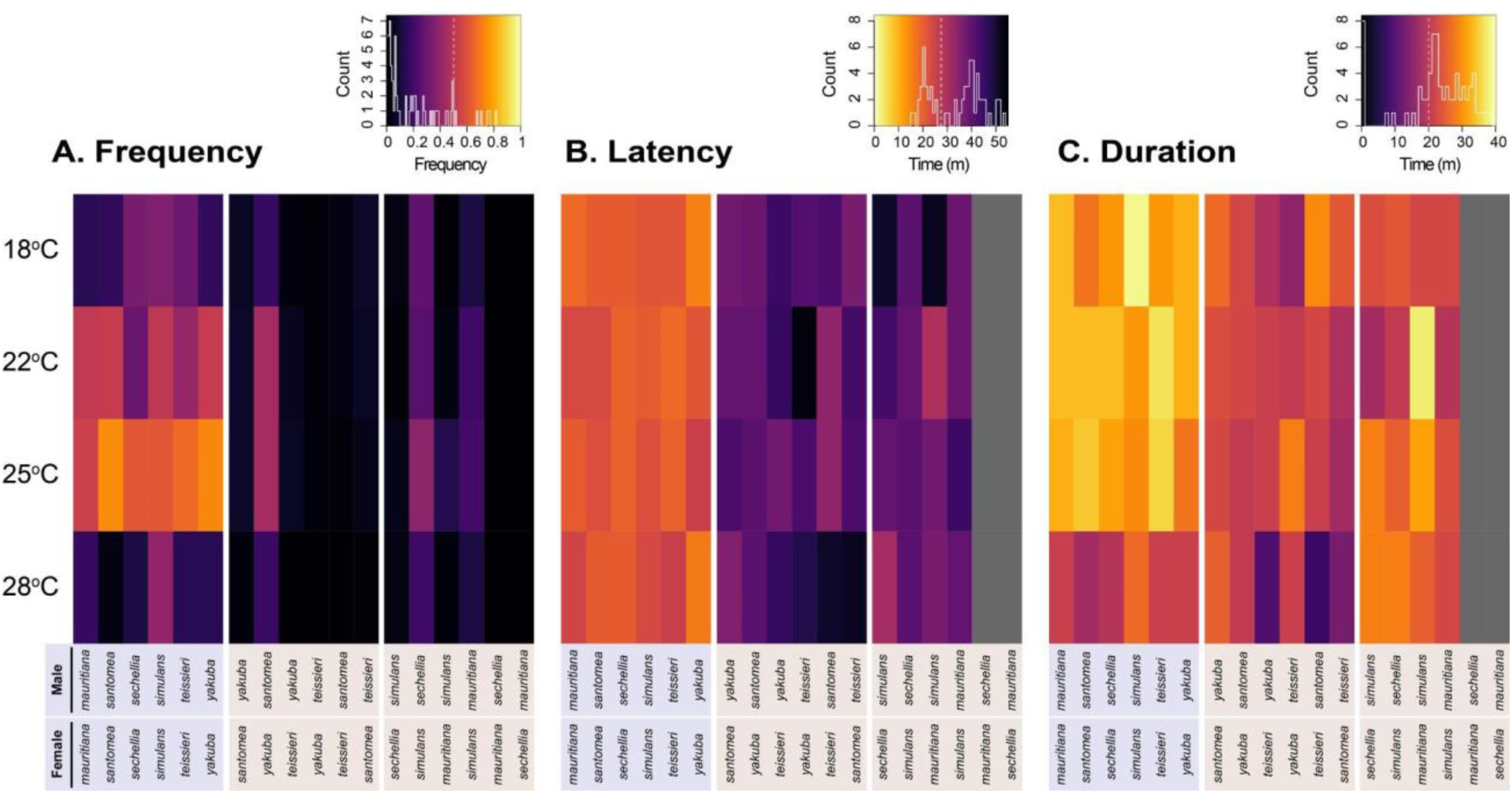
Non-choice individual mating trials reveal a small effect of temperature in heterospecific mating propensity. A subset of the conspecific crosses shown in Figure 2 is shown for comparison. **A.** Mating frequency measured as a proportion of successful matings over total trials for two *Drosophila* species triads. **B.** Mating latency measured in minutes. **C.** Mating duration measured in minutes.

**FIGURE 6.**
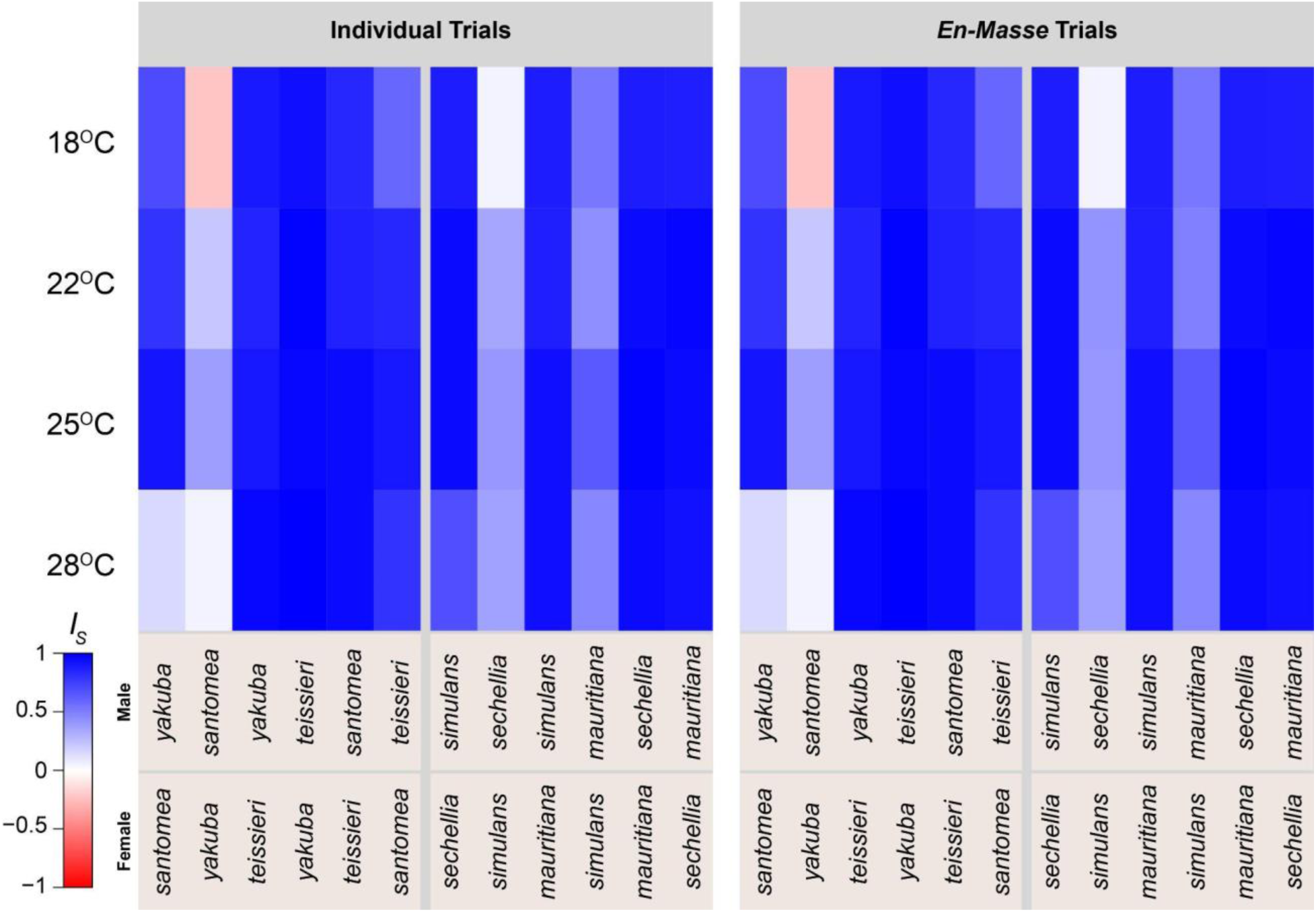
*I_S_*, a metric of sexual isolation between species, is dependent on temperature in some interspecific crosses. The reciprocal crosses between *D. yakuba* and *D. santomea* showed instances of lower-than 0, *I_S_*, which is caused by heterospecific and conspecific being equally likely at a given temperature.

## DISCUSSION

In this report, we describe the effect of temperature on conspecific-mating frequency and behavior in fourteen species of *Drosophila*, and find that while temperature is a determinant of the likelihood of mating, the effect is highly species-specific. We also find that the effect of temperature on the frequency of heterospecific matings in six interspecific hybridizations is moderate. These results have three implications. First, they indicate that mating propensity is dependent upon environmental temperature in *Drosophila*. Since flies use external temperature to thermoregulate, it follows that flies should be more receptive in their optimal temperature range, and that they will be less receptive at stressful temperatures. Second, our results suggest that environmental temperature might influence hybridization rates between species not by increasing the likelihood of heterospecific matings, but rather by reducing the likelihood of conspecific matings. Collectively, these results indicate that conditions in which mating takes place are an overlooked component of the experimental design of behavioral mating experiments in animals.

Our study follows in the steps of previous research that has suggested that while temperature can affect mating behaviors, that effect is variable. A meta-analysis on the effect of within-species mating behavior across animals suggested no consistent directional effect of temperature on mating behaviors and mating success (Pilakouta and Baillet, 2022). These observations are consistent with our results and both studies highlight the difficulties of predicting changes in the strength of sexual selection among natural populations in a warming world (Pilakouta and Ålund 2021). One potential mechanism for the differences in mating likelihood is that individuals are more physiologically stressed in maintaining homeostasis at higher temperatures, and that they are simply not active at lower temperatures. Other possibilities also exist. Temperature might affect mating by modulating the transduction or reception of courtship signals. Rearing temperature affects cuticular hydrocarbon profiles, which can be important in mate choice across insects (Conrad et al., 2017; Duarte et al., 2019; Kárpáti et al., 2023; Michelutti et al., 2018; Noorman and Otter, 2002; Rajpurohit et al., 2021; Savarit and Ferveur, 2002; Silva et al. 2007; Westerman and Monteiro 2016). Mating temperature affects visual and vibratory courtship behaviors in spiders, (*Habronattus clypeatus*, Brandt et al., 2020, 2018; *Schizocosa floridana*, Rosenthal and Elias, 2019). In rock lizards (*Iberolacerta cyreni*), chemosensory signals are less likely to be effectively conducted at high temperatures (Martín and López, 2013). Dissecting the precise molecular, physiological, and neurological underpinnings of the interaction between mating choice and temperature will be of critical importance to understand species boundaries in the face of global warming (Chunco, 2014; Groot and Zizzari, 2019; Muhlfeld et al., 2014; Vallejo-Marín and Hiscock, 2016).

The second implication of our findings is the effect of temperature on the likelihood of heterospecific matings. Our findings indicate that while the rate of conspecific matings is highly contingent on environmental temperature, the rate of interspecific matings is roughly similar across temperatures. These results imply that environmental conditions can play a key role in the proportional representation of heterospecific vs. conspecific matings, and therefore in the establishment and persistence of hybrid zones. It has been suggested that instances of hybridization might be more likely to occur at the edge of a geographic range because of the relative rarity of conspecifics (and abundance of heterospecifics, Wilson and Hedrick, 1982), but hybrid zones also tend to occur at the edge of species limits where physiological boundaries are pushed. In the case of *Drosophila*, *D. santomea* and *D. yakuba* hybridize in the midlands of the oceanic volcano of São Tomé (Lachaise et al., 2000; Llopart et al., 2005; Matute, 2010) where the hybrid zone occurs at the upper edge of the thermal range for *D. santomea*, and at the lower edge for *D. yakuba*. The competitive outcomes between these two species are mediated by temperature, with *D. yakuba* outperforming *D. santomea* in warmer conditions (Comeault and Matute, 2021). Similarly, *D. teissieri* and *D. yakuba* hybridize in the highlands of another island, Bioko, at the lower end of the thermal range of *D. teissieri* (Cooper et al., 2018). Our experiments here reflect the importance of thermal fitness differences in these species pairs and demonstrate that the likelihood of mating in both *D. santomea* and *D. teissieri* —the thermally sensitive species—is affected more strongly by temperature than that of *D. yakuba*.

Other environmental conditions, besides temperature, are also important modulators of mating behavior and reproductive isolation. Temperature, density, and age are all environmental factors that affect the extent of phenotypic variation in traits that lead to behavioral isolation between nascent and well-formed species, alike. Both modeling (Reeve, 1989; Shizuka and Hudson, 2020; Wilson and Hedrick, 1982) and experimental studies (Friberg et al., 2013; Gomez-Llano et al., 2018; Keränen et al., 2013; Matute, 2014) have shown that the density of heterospecifics is an important factor in mating propensity. Speciation can be impeded by mating choice, as individuals that have few choices might pursue matings with heterospecifics, even if the offspring are somehow less fit, depending on the waiting time for a potential conspecific mate (Chen and Pfennig, 2020; Wilson and Hedrick, 1982). This may be especially pronounced in species that have low abundance at the edge of their range, and which have the potential to hybridize at that interface. A third component that might determine the likelihood of heterospecific matings is the age of the individuals engaging in the cross. In *Aedes* mosquitos, older individuals tend to engage in heterospecific matings more readily than younger ones (Bargielowski et al., 2019). The density of conspecifics and heterospecifics, their age, and environmental temperature might all covary and their relative effect in reproductive isolation might be challenging to dissect in field conditions.

Besides sexual isolation, developmentally-based reproductive isolation is also affected by environmental conditions, and genetic studies provide a mechanism for this genetic × environment interaction. A handful of studies have shown that hybrid inviability and hybrid sterility are both dependent on temperature (Coyne et al., 1998; Lee 1978; Mason et al., 2011; Presgraves, 2003; Wongpatsa et al., 2014). Hybrids between *Drosophila* species show higher levels of inviability at higher temperatures (Barbash et al., 2000; Hutter and Ashburner, 1987; Lee, 1978; Matute et al., 2010; Sawamura et al., 1993), and genome-wide mapping suggests that this effect is generalized across the genome. In *melanogaster/santomea* and *melanogaster/simulans* hybrids, the penetrance of different genomic regions involved in inviability is greater at higher temperatures than at low temperatures (Coyne et al., 1998; Matute et al., 2010; Miller and Matute, 2017; Presgraves, 2003). Several genetic loci involved in hybrid fitness reductions at 24°C have diminished effects at lower temperatures. In hybrids between *Tribolium* beetle species, hybrid male viability declines as temperature increases, leading to increasing manifestation of Haldane’s rule at high temperatures (Wade et al., 1999). These previous results suggest that reproductive isolation even between long-diverged species is conditioned by temperature. The effect of age is not limited to premating isolation and can also affect the strength of postzygotic isolation. In hybrids between the subspecies of *D. pseudoobscura*, hybrid males are weakly fertile but only when they are old (Orr and Irving, 2005). These instances suggest that even for barriers that are often thought to be intrinsic, ecological factors might be of critical importance (Anderson et al., 2023).

Our study also has some important caveats. The flies in our study (both male and female) were kept in temperature-controlled conditions throughout the trials and they were reared in the same temperature regimes at which they were tested. We are therefore unable to comment on any influence on mating preferences among these species that might arise from development in other temperature conditions or in fluctuating conditions. Reproductive traits like fecundity and fertility are also dependent on developmental temperature in *Drosophila* (Huey et al. 1995; Nuney and Cheung 1997; Matute et al. 2009; Kelpsatel et al. 2019; Comeault et al. 2020) and other insects (Kersting et al. 1999, Papanikolaou et al. 2013, Cui et al. 2018, Li et al. 2020). Understanding how thermal regimes during development interact with climate exposure as reproductive adults work to shape mating choices thus merits follow-up. Second, our experiments are necessarily unrealistic. *En-masse* experiments tested female choice when presented with multiple males simultaneously generating male-male, and female-female interactions, which our experimental design does not allow us to quantify. Similarly, individual non-choice experiments include only one female, and one male, which might represent an oversimplification of natural matings. It remains possible that females might react differently when presented with males in a more sequential fashion, and that the order in which a female encounters con-vs. heterospecific males might alter her mating preferences or other aspects of her physiology (e.g., Matute and Coyne 2010).

Our report serves as a survey demonstrating that behavioral traits are affected by environmental conditions. While this is not a particularly surprising result, interactions between environmental factors and the alleles that underlie behavior and reproductive isolation remain largely understudied. In order to truly understand how mating choice and reproductive isolation occur in nature, they need to be studied in the context of the conditions in which they take place.

## Acknowledgements

We would like to thank the Matute lab for helpful comments. JAR was supported by the National Institute of Environmental Health Sciences under award F32ES035271. DRM was supported by the National Institute of General Medical Sciences under Award R35GM148244.

## Supplemental Material

### Supplementary Figures

**FIGURE S1.**
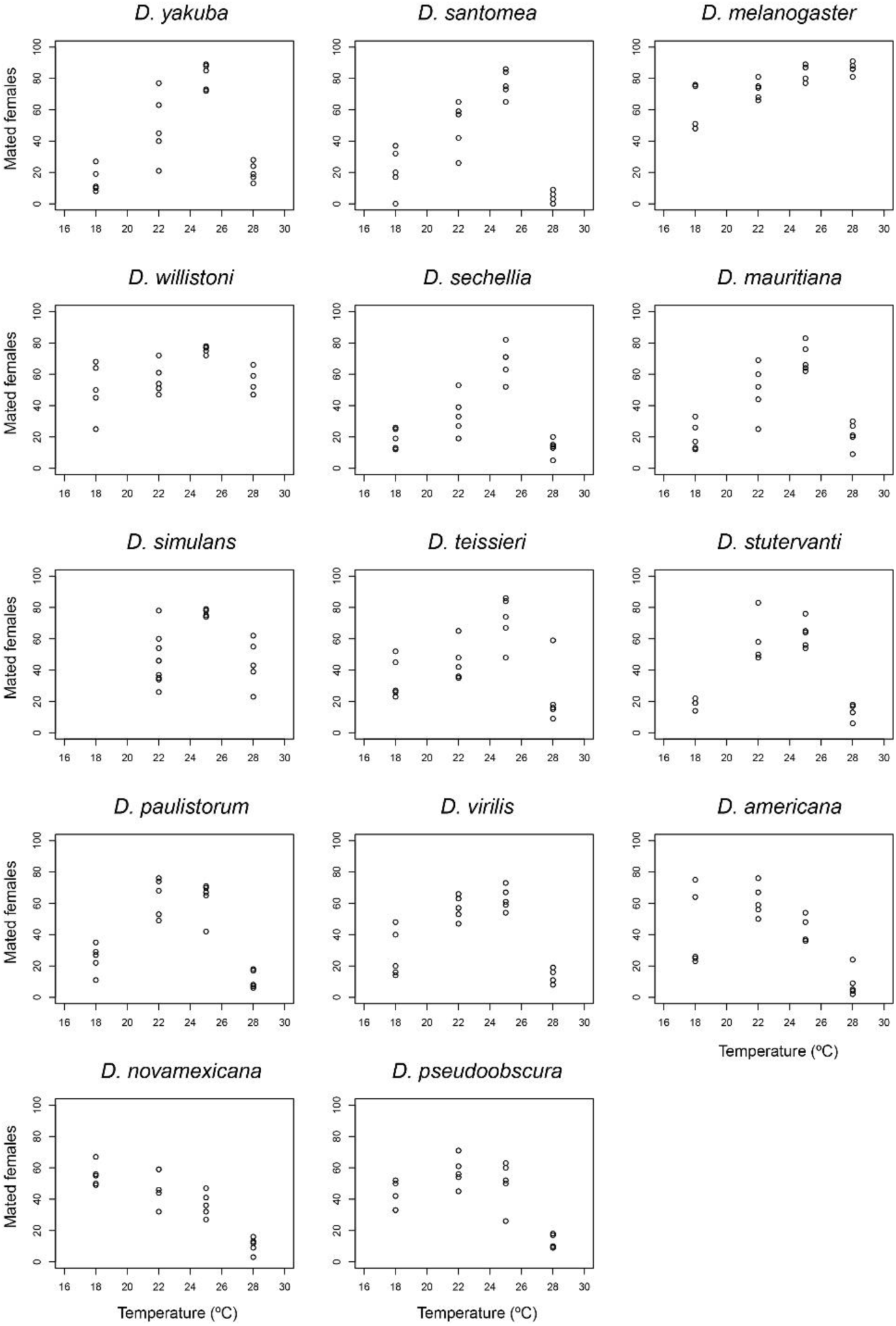
Proportion of females mated for each of the fourteen species included in this study. Note that unlike the linear models we present in the text which had a binomial response (mated vs. unmated), each point in these panels show the proportion of mated females in a replicate.

**FIGURE S2.**
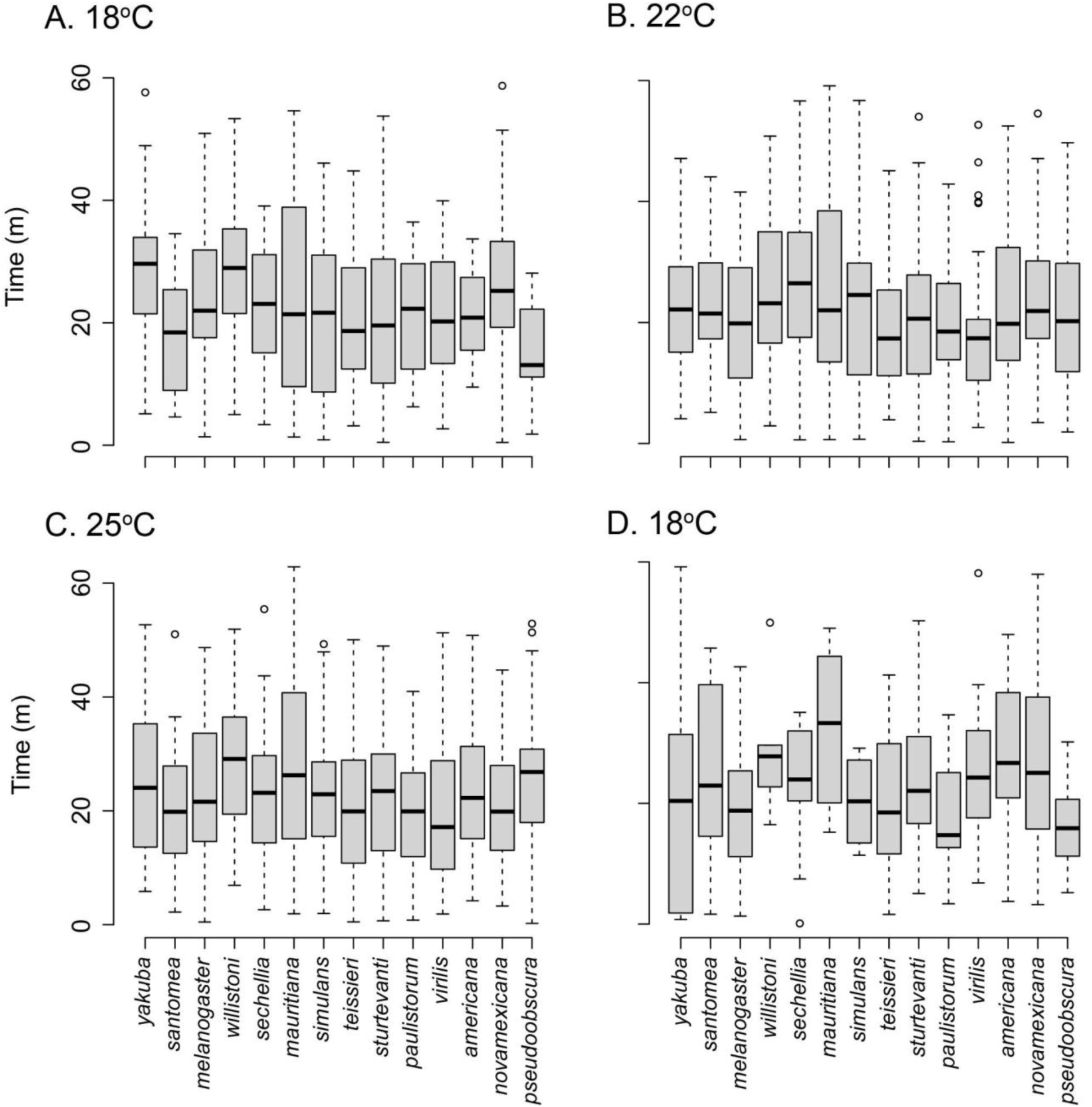
Conspecific mating latency during the individual experiments at four temperatures.

**FIGURE S3.**
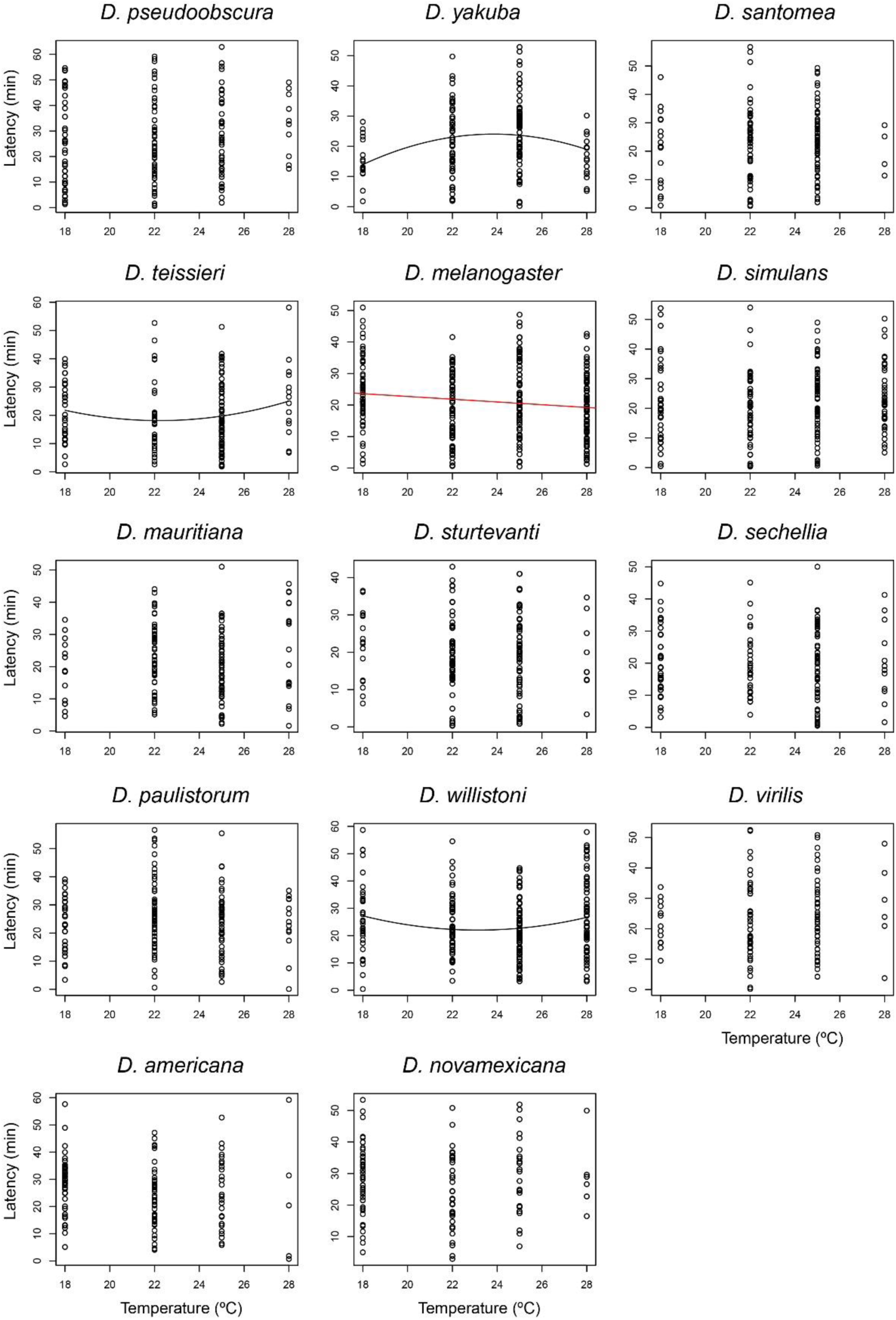
Conspecific mating latency shown for each species along the assayed temperature range.

**FIGURE S4.**
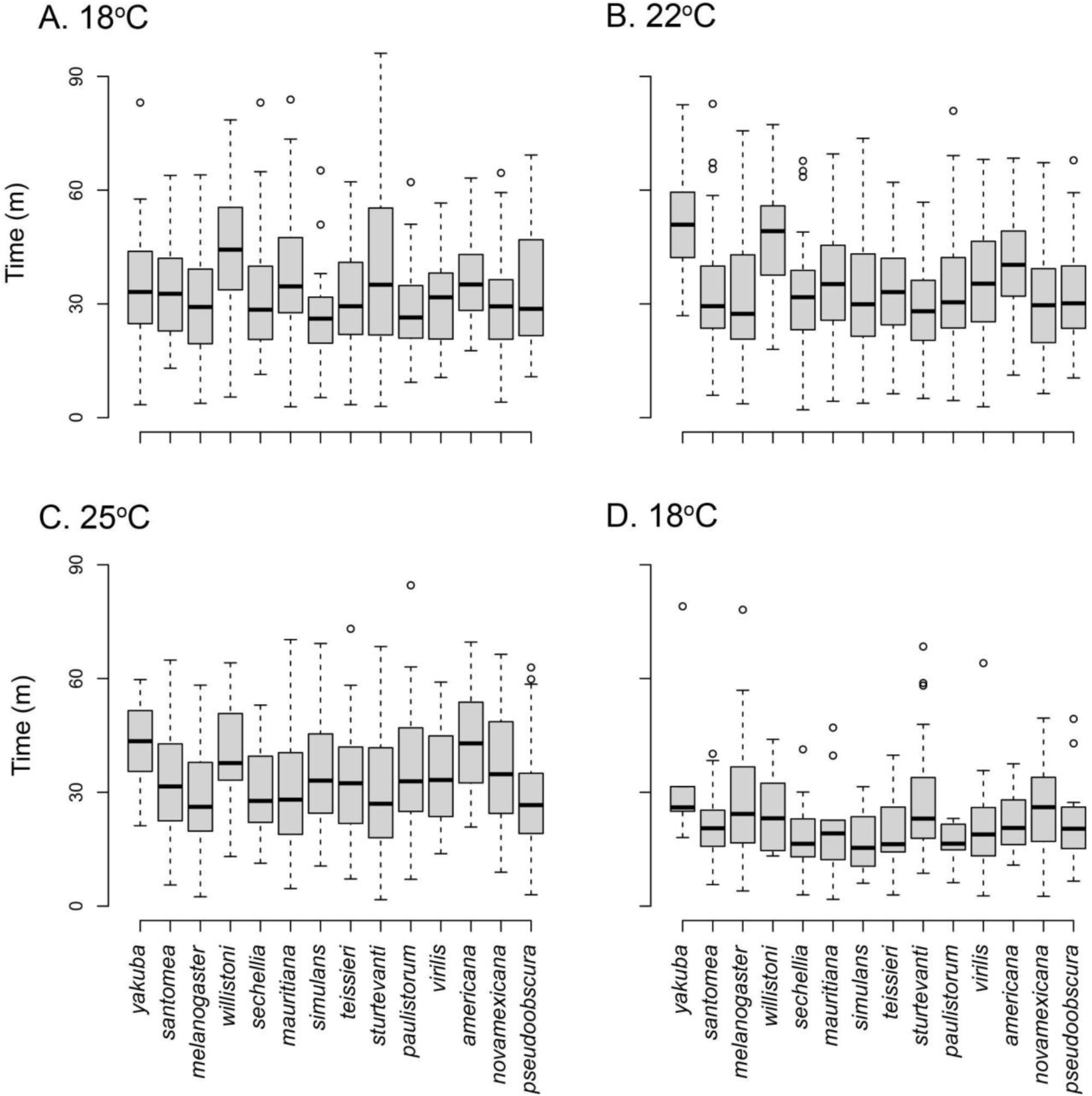
Conspecific mating duration during the individual experiments at four temperatures.

**FIGURE S5.**
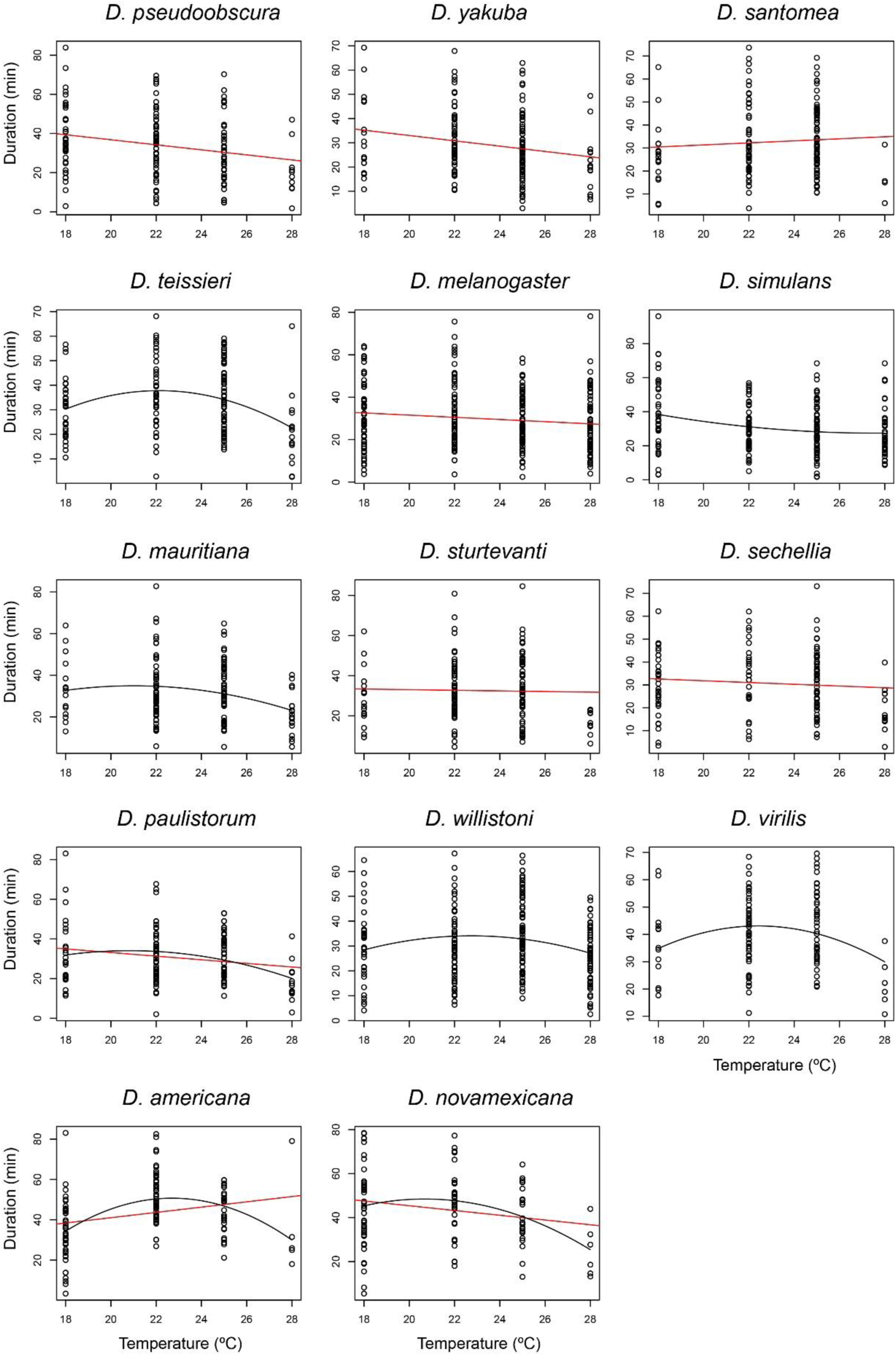
Conspecific mating duration shown for each species along the assayed temperature range.

### Supplementary Tables

**TABLE S1.** Isofemale lines used in this study.

| **Species** | **Isofemale line** | **Collector** | **Location** | **Year** |
| --- | --- | --- | --- | --- |
| *D. sechellia* | Anro105 | Matute, D.R. | Anse Royale, La Digue, Seychelles | 2012 |
| *D. simulans* | MP2.45 | Matute, D.R. | Mpala, Kenya | 2018 |
| *D. santomea* | TNE1500.2 | Matute, D.R. | Bom sucesso, São Tomé é Príncipe | 2015 |
| *D. yakuba* | BOSU1151 | Matute, D.R. | San Nicolau, São Tomé é Príncipe | 2015 |
| *D. teissieri* | NM5.1 | Matute, D.R. | Bioko, Equatorial Guinea | 2009 |
| *D. melanogaster* | Loreto | Matute, D.R. | Bioko, Equatorial Guinea | 2009 |
| *D. mauritiana* | Standard | Kitagawa, O. | Mauritius | 1981 |
| *D. pseudoobscura* | 14011-0121.117 | Mateos, M. | Tucson, Arizona | 2004 |
| *D. novamexicana* | 15010-1031.14 | Castrezana, S. | Moab, Utah | 1949 |
| *D. americana* | 15010-0951.22 | McAllister, B. | Illinois River at Duck Island, Illinois | 2005 |
| *D. virilis* | 15010-1051.00 | NA | Pasadena, California. | Before 1984 |
| *D. willistoni* | 14030-0811.32 | Bruck, J. | Monkey Hill, St. Kitts, | 2005 |
| *D. paulistorum* | 14030-0771.00 | NA | Copan, Honduras. | Before 1984 |
| *D. sturtevantii* | 14043-0871.01 | Heed, W. | Bucaramanga, Colombia | 1956 |

**TABLE S2.** Interspecific hybridizations included in this study.

| **Mother** | **Father** |
| --- | --- |
| *D. yakuba* | *D. santomea* |
| *D. santomea* | *D. yakuba* |
| *D. yakuba* | *D. teissieri* |
| *D. teissieri* | *D. yakuba* |
| *D. santomea* | *D. teissieri* |
| *D. teissieri* | *D. yakuba* |
| *D. simulans* | *D. sechellia* |
| *D. sechellia* | *D. simulans* |
| *D. sechellia* | *D. mauritiana* |
| *D. sechellia* | *D. sechellia* |
| *D. mauritiana* | *D. simulans* |
| *D. simulans* | *D. mauritiana* |

**TABLE S3.** Akaike Information Criterion (AIC) values for linear logistic and quadratic logistic models to study the effect of temperature and cross direction in heterospecific mating frequency in *en-masse* mating experiments.

| Species pair | AIC<sub>Linear</sub> | AIC<sub>Quadratic</sub> |
| --- | --- | --- |
| *D. yakuba/D. santomea* | 3,453.98 | 3,297.38 |
| *D. yakuba/D. teissieri* | 968.34 | 941.56 |
| *D. santomea/D. teissieri* | 1,524.44 | 1,505.53 |
| *D. simulans/D. sechellia* | 3,089.12 | 3,083.27 |
| *D. simulans/D. mauritiana* | 2,837.22 | 2,809.49 |
| *D. sechellia/D. mauritiana* | 698.25 | 702.11 |

**TABLE S4.** Linear logistic models suggest that temperature has a moderate effect on mating propensity in six types of heterospecific *en-masse* matings. All the likelihood ratio tests (LRT) comparisons involve one degree of freedom.

| Species pair | Direction |  | Temperature |  | Direction x temperature |  |
| --- | --- | --- | --- | --- | --- | --- |
| | $\chi^2$ | *P* | $\chi^2$ | *P* | $\chi^2$ | *P* |
| *D. yakuba*/<br>*D. santomea* | 437.567 | < 1 ×10<sup>-10</sup> | 1.0658 | 0.302 | **9.120** | **0.003** |
| *D. yakuba*/<br>*D. teissieri* | 43.888 | < 1 ×10<sup>-10</sup> | 2.396 | 0.122 | 0.187 | 0.666 |
| *D. santomea*/<br>*D. teissieri* | 37.072 | 1.139×10<sup>-9</sup> | **11.175** | **8.290×10<sup>-4</sup>** | 3.039<br>×10<sup>-4</sup> | 0.986 |
| *D. simulans*/<br>*D. sechellia* | 779.092 | < 1 ×10<sup>-10</sup> | 0.167 | 0.683 | **7.862** | **0.005** |
| *D. simulans*/<br>*D. mauritiana* | 388.847 | < 1 ×10<sup>-10</sup> | 2.495 | 0.114 | 2.780 | 0.094 |
| *D. sechellia*/<br>*D. mauritiana* | 0.118 | 0.732 | **4.067** | **0.044** | 0.687 | 0.407 |

**TABLE S5.** Correlation test of mating frequencies between *en-masse* and individual non-choice experiments.

| Temperature (°C) | Pearson's product-moment correlation | *P*-value |
| --- | --- | --- |
| 18 | 0.848 | 1.281 × 10<sup>-4</sup> |
| 22 | 0.749 | 2.044 × 10<sup>-3</sup> |
| 25 | 0.953 | 1.497 × 10<sup>-7</sup> |
| 28 | 0.987 | 7.783 × 10<sup>-11</sup> |

**TABLE S6.** AIC values and effect significance (LRT results) for linear logistic and quadratic logistic models to study the effect of temperature and cross direction in heterospecific mating frequency in individual non-choice mating experiments.

| Species | Quadratic Model |  |  | Linear Model | AIC |
| --- | --- | --- | --- | --- | --- |
| | $l(\text{temperature}^2)$ | Temperature | AIC | Temperature | |
| *D. pseudoobscura* | $\chi^2_1 = 28.419$<br>$P = 9.773\text{e}^{-08}$ | $\chi^2_1 = 24.942$<br>$P = 5.907\text{e}^{-07}$ | 509.9356 | $\chi^2_1 = 21.541$<br>$P = 3.463\text{e}^{-06}$ | **508.5483** |
| *D. yakuba* | $\chi^2_1 = 111.15$<br>$P < 2.2\text{e}^{-16}$ | $\chi^2_1 = 112.64$<br>$P < 1 \times 10^{-10}$ | **471.1564** | $\chi^2_1 = 1.5657$<br>$P = 0.2108$ | 540.0222 |
| *D. santomea* | $\chi^2_1 = 160.45$<br>$P < 2.2\text{e}^{-16}$ | $\chi^2_1 = 159.45$<br>$P < 1 \times 10^{-10}$ | **425.9435** | $\chi^2_1 = 0.021533$<br>$P = 0.8833$ | 532.1967 |
| *D. teissieri* | $\chi^2_1 = 39.629$<br>$P = 3.071\text{e}^{-10}$ | $\chi^2_1 = 38.839$<br>$P = 4.602\text{e}^{-10}$ | **528.3322** | $\chi^2_1 = 0.50133$<br>$P = 0.4789$ | 536.6658 |
| *D. melanogaster* | $\chi^2_1 = 0.159706$<br>$P = 0.6894$ | $\chi^2_1 = 0.018713$<br>$P = 0.8912$ | **425.9435** | $\chi^2_1 = 15.165$<br>$P = 9.854\text{e}^{-05}$ | 532.1967 |
| *D. simulans* | $\chi^2_1 = 12.561$<br>$P = 0.0003938$ | $\chi^2_1 = 13.102$<br>$P = 0.0002949$ | 573.4243 | $\chi^2_1 = 1.5361$<br>$P = 0.2152$ | **555.771** |
| *D. mauritiana* | $\chi^2_1 = 68.146$<br>$P < 2.2\text{e}^{-16}$ | $\chi^2_1 = 69.123$<br>$P < 1 \times 10^{-10}$ | **486.3818** | $\chi^2_1 = 1.1626$<br>$P = 0.2809$ | 520.7947 |
| *D. sturtevantii* | $\chi^2_1 = 106.33$<br>$P < 2.2\text{e}^{-16}$ | $\chi^2_1 = 105.55$<br>$P < 1 \times 10^{-10}$ | **457.2496** | $\chi^2_1 = 0.042049$<br>$P = 0.8375$ | 523.1424 |
| *D. sechellia* | $\chi^2_1 = 20.611$<br>$P = 5.626\text{e}^{-06}$ | $\chi^2_1 = 19.646$<br>$P = 9.321\text{e}^{-06}$ | 525.9057 | $\chi^2_1 = 2.0594$<br>$P = 0.1513$ | **517.3777** |
| *D. paulistorum* | $\chi^2_1 = 42.731$<br>$P = 6.280\text{e}^{-11}$ | $\chi^2_1 = 40.804$<br>$P = 1.683\text{e}^{-10}$ | **506.288** | $\chi^2_1 = 3.711$<br>$P = 0.05405$ | 518.2463 |
| *D. willistoni* | $\chi^2_1 = 8.4439$<br>$P = 0.003663$ | $\chi^2_1 = 10.1216$<br>$P = 0.001465$ | **425.9435** | $\chi^2_1 = 17.224$<br>$P = 3.321\text{e}^{-05}$ | 532.1967 |
| *D. virilis* | $\chi^2_1 = 74.908$<br>$P < 2.2\text{e}^{-16}$ | $\chi^2_1 = 73.927$<br>$P < 1 \times 10^{-10}$ | **426.4345** | $\chi^2_1 = 0.27382$<br>$P = 0.6008$ | 472.3098 |
| *D. americana* | $\chi^2_1 = 11.7211$<br>$P = 0.000618$ | $\chi^2_1 = 8.8163$<br>$P = 0.002986$ | 470.7596 | $\chi^2_1 = 40.809$<br>$P = 1.679\text{e}^{-10}$ | **456.86** |
| *D. novamexicana* | $\chi^2_1 = 2.1733$<br>$P = 0.1404$ | $\chi^2_1 = 1.1047$<br>$P = 0.2932$ | 450.7751 | $\chi^2_1 = 37.175$<br>$P = 1.08\text{e}^{-09}$ | **431.4303** |

**TABLE S7.**
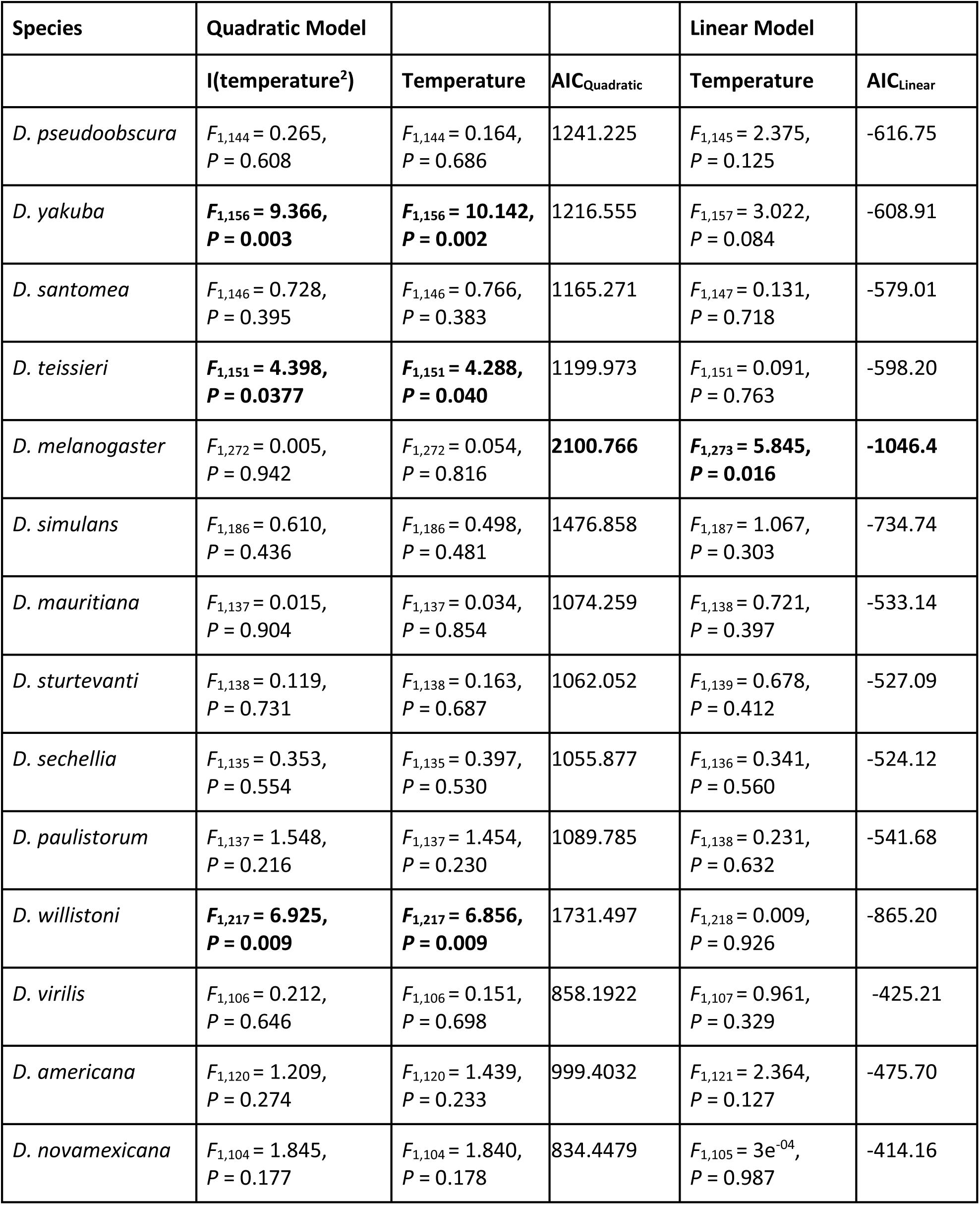
Effect of temperature on mating latency in conspecific crosses for 14 different species of *Drosophila*.

| Species | Quadratic Model |  |  | Linear Model |  |
| --- | --- | --- | --- | --- | --- |
|  | I(temperature<sup>2</sup>) | Temperature | AIC<sub>Quadratic</sub> | Temperature | AIC<sub>Linear</sub> |
| *D. pseudoobscura* | $F_{1,144} = 0.265$,<br>$P = 0.608$ | $F_{1,144} = 0.164$,<br>$P = 0.686$ | 1241.225 | $F_{1,145} = 2.375$,<br>$P = 0.125$ | -616.75 |
| *D. yakuba* | **$F_{1,156} = 9.366$,**<br>**$P = 0.003$** | **$F_{1,156} = 10.142$,**<br>**$P = 0.002$** | 1216.555 | $F_{1,157} = 3.022$,<br>$P = 0.084$ | -608.91 |
| *D. santomea* | $F_{1,146} = 0.728$,<br>$P = 0.395$ | $F_{1,146} = 0.766$,<br>$P = 0.383$ | 1165.271 | $F_{1,147} = 0.131$,<br>$P = 0.718$ | -579.01 |
| *D. teissieri* | **$F_{1,151} = 4.398$,**<br>**$P = 0.0377$** | **$F_{1,151} = 4.288$,**<br>**$P = 0.040$** | 1199.973 | $F_{1,151} = 0.091$,<br>$P = 0.763$ | -598.20 |
| *D. melanogaster* | $F_{1,272} = 0.005$,<br>$P = 0.942$ | $F_{1,272} = 0.054$,<br>$P = 0.816$ | **2100.766** | **$F_{1,273} = 5.845$,**<br>**$P = 0.016$** | **-1046.4** |
| *D. simulans* | $F_{1,186} = 0.610$,<br>$P = 0.436$ | $F_{1,186} = 0.498$,<br>$P = 0.481$ | 1476.858 | $F_{1,187} = 1.067$,<br>$P = 0.303$ | -734.74 |
| *D. mauritiana* | $F_{1,137} = 0.015$,<br>$P = 0.904$ | $F_{1,137} = 0.034$,<br>$P = 0.854$ | 1074.259 | $F_{1,138} = 0.721$,<br>$P = 0.397$ | -533.14 |
| *D. sturtevantii* | $F_{1,138} = 0.119$,<br>$P = 0.731$ | $F_{1,138} = 0.163$,<br>$P = 0.687$ | 1062.052 | $F_{1,139} = 0.678$,<br>$P = 0.412$ | -527.09 |
| *D. sechellia* | $F_{1,135} = 0.353$,<br>$P = 0.554$ | $F_{1,135} = 0.397$,<br>$P = 0.530$ | 1055.877 | $F_{1,136} = 0.341$,<br>$P = 0.560$ | -524.12 |
| *D. paulistorum* | $F_{1,137} = 1.548$,<br>$P = 0.216$ | $F_{1,137} = 1.454$,<br>$P = 0.230$ | 1089.785 | $F_{1,138} = 0.231$,<br>$P = 0.632$ | -541.68 |
| *D. willistoni* | **$F_{1,217} = 6.925$,**<br>**$P = 0.009$** | **$F_{1,217} = 6.856$,**<br>**$P = 0.009$** | 1731.497 | $F_{1,218} = 0.009$,<br>$P = 0.926$ | -865.20 |
| *D. virilis* | $F_{1,106} = 0.212$,<br>$P = 0.646$ | $F_{1,106} = 0.151$,<br>$P = 0.698$ | 858.1922 | $F_{1,107} = 0.961$,<br>$P = 0.329$ | -425.21 |
| *D. americana* | $F_{1,120} = 1.209$,<br>$P = 0.274$ | $F_{1,120} = 1.439$,<br>$P = 0.233$ | 999.4032 | $F_{1,121} = 2.364$,<br>$P = 0.127$ | -475.70 |
| *D. novamexicana* | $F_{1,104} = 1.845$,<br>$P = 0.177$ | $F_{1,104} = 1.840$,<br>$P = 0.178$ | 834.4479 | $F_{1,105} = 3e^{-04}$,<br>$P = 0.987$ | -414.16 |

**TABLE S8.** Linear and quadratic logistic models suggest that temperature has a moderate effect in the duration of some conspecific matings.

| Species | Quadratic Model |  | AIC<sub>Quadratic</sub> | Linear Model | AIC<sub>Linear</sub> |
| --- | --- | --- | --- | --- | --- |
|  | **l(temperature<sup>2</sup>)** | **Temperature** |  | **Temperature** |  |
| *D. pseudoobscura* | $F_{1,144} = 1.826$,<br>$P = 0.179$ | $F_{1,156} = 1.312$,<br>$P = 0.254$ | **1249.029** | $F_{1,144} = \mathbf{8.215}$,<br>$P = \mathbf{0.005}$ | 1248.882 |
| *D. yakuba* | $F_{1,156} = 1.546$,<br>$P = 0.2156$ | $F_{1,156} = 1.082$,<br>$P = 0.300$ | **1285.06** | $F_{1,157} = \mathbf{7.190}$,<br>$P = \mathbf{0.008}$ | 1285.492 |
| *D. santomea* | $F_{1,146} = \mathbf{4.739}$,<br>$P = \mathbf{0.031}$ | $F_{1,146} = \mathbf{4.972}$,<br>$P = \mathbf{0.027}$ | 1230.471 | $F_{1,147} = 0.763$,<br>$P = 0.384$ | 1233.23 |
| *D. teissieri* | $F_{1,151} = \mathbf{14.875}$,<br>$P = \mathbf{0.0002}$ | $F_{1,151} = \mathbf{14.399}$,<br>$P = \mathbf{0.0002}$ | 1246.852 | $F_{1,152} = 0.531$,<br>$P = 0.467$ | 1259.321 |
| *D. melanogaster* | $F_{1,272} = 0.555$,<br>$P = 0.457$ | $F_{1,272} = 0.360$,<br>$P = 0.549$ | 2251.452 | $F_{1,273} = \mathbf{4.664}$,<br>$P = \mathbf{0.032}$ | **2250.012** |
| *D. simulans* | $F_{1,186} = 1.282$,<br>$P = 0.259$ | $F_{1,186} = 1.845$,<br>$P = 0.176$ | 1586.757 | $F_{1,187} = \mathbf{10.406}$,<br>$P = \mathbf{0.001}$ | **1586.055** |
| *D. mauritiana* | $F_{1,137} = 3.835$,<br>$P = 0.052$ | $F_{1,137} = 3.159$,<br>$P = 0.078$ | **1143.715** | $F_{1,138} = \mathbf{5.290}$,<br>$P = \mathbf{0.023}$ | 1145.58 |
| *D. sturtevantii* | $F_{1,138} = \mathbf{7.997}$,<br>$P = \mathbf{0.005}$ | $F_{1,138} = \mathbf{7.827}$,<br>$P = \mathbf{0.006}$ | 1173.91 | $F_{1,139} = 0.092$,<br>$P = 0.763$ | 1179.853 |
| *D. sechellia* | $F_{1,135} = \mathbf{7.918}$,<br>$P = \mathbf{0.006}$ | $F_{1,135} = \mathbf{7.498}$,<br>$P = \mathbf{0.007}$ | 1112.746 | $F_{1,136} = 1.107$,<br>$P = 0.295$ | 1118.611 |
| *D. paulistorum* | $F_{1,137} = \mathbf{5.894}$,<br>$P = \mathbf{0.016}$ | $F_{1,137} = \mathbf{5.019}$,<br>$P = \mathbf{0.027}$ | **1122.547** | $F_{1,138} = \mathbf{5.885}$,<br>$P = \mathbf{0.017}$ | 1126.445 |
| *D. willistoni* | $F_{1,217} = \mathbf{8.676}$,<br>$P = \mathbf{0.004}$ | $F_{1,217} = \mathbf{8.294}$,<br>$P = \mathbf{0.004}$ | 1790.954 | $F_{1,218} = 0.666$,<br>$P = 0.415$ | 1797.579 |
| *D. virilis* | $F_{1,106} = \mathbf{7.293}$,<br>$P = \mathbf{0.008}$ | $F_{1,106} = \mathbf{7.169}$,<br>$P = \mathbf{0.009}$ | 883.6012 | $F_{1,107} = 0.045$,<br>$P = 0.832$ | 888.8541 |
| *D. americana* | $F_{1,120} = \mathbf{27.008}$,<br>$P = \mathbf{8.419e-07}$ | $F_{1,120} = \mathbf{29.115}$,<br>$P = \mathbf{3.485e-07}$ | **999.4032** | $F_{1,121} = \mathbf{8.218}$,<br>$P = \mathbf{0.005}$ | 1022.371 |
| *D. novamexicana* | $F_{1,104} = \mathbf{6.281}$,<br>$P = \mathbf{0.014}$ | $F_{1,104} = \mathbf{5.558}$,<br>$P = \mathbf{0.020}$ | **905.669** | $F_{1,105} = \mathbf{4.546}$,<br>$P = \mathbf{0.035}$ | 909.9437 |

**TABLE S9.** AIC values for heterospecific linear logistic and quadratic logistic models using the mating frequency in heterospecific individual non-choice matings.

| Species pair | AIC<sub>Linear</sub> | AIC<sub>Quadratic</sub> |
| --- | --- | --- |
| *D. yakuba* / *D. santomea* | **699.080** | **668.577** |
| *D. yakuba* / *D. teissieri* | 226.821 | 225.142 |
| *D. santomea* / *D. teissieri* | 273.291 | 275.879 |
| *D. simulans* / *D. sechellia* | 609.200 | 609.775 |
| *D. simulans* / *D. mauritiana* | **537.200** | **528.301** |
| *D. sechellia* / *D. mauritiana* | NA | NA |

**TABLE S10.** Linear logistic models suggest that temperature has a moderate effect on mating propensity in some of the six types of heterospecific *en-masse* matings. The metric of isolation is receptivity of females in *en-masse* matings. All the likelihood ratio tests (LRT) comparisons involve one degree of freedom.

| Species pair | Direction |  | Temperature |  | Direction × temperature |  |
| --- | --- | --- | --- | --- | --- | --- |
| | $\chi^2$ | *P* | $\chi^2$ | *P* | $\chi^2$ | *P* |
| *D. yakuba*/<br>*D. santomea* | **88.465** | **< 1 × 10<sup>-10</sup>** | 4.114×10<sup>-5</sup> | 0.995 | 1.304 | 0.254 |
| *D. yakuba*/<br>*D. teissieri* | 3.420 | 0.064 | 0.258 | 0.611 | 7.882×10<sup>-7</sup> | 0.999 |
| *D. santomea*/<br>*D. teissieri* | **82.65** | **< 1 × 10<sup>-10</sup>** | **5.843** | **0.016** | 4.961×10<sup>-3</sup> | 0.944 |
| *D. simulans*/<br>*D. sechellia* | **98.171** | **< 1 × 10<sup>-10</sup>** | 0.183 | 0.669 | 0.260 | 0.610 |
| *D. simulans*/<br>*D. mauritiana* | **23.387** | **1.324 × 10<sup>-6</sup>** | 0.812 | 0.368 | 1.098 | 0.295 |
| *D. sechellia*/<br>*D. mauritiana* | 0.58! | 0.446 | **3.9648** | **0.04646** | 0.687 | 0.407 |

**TABLE S11.**
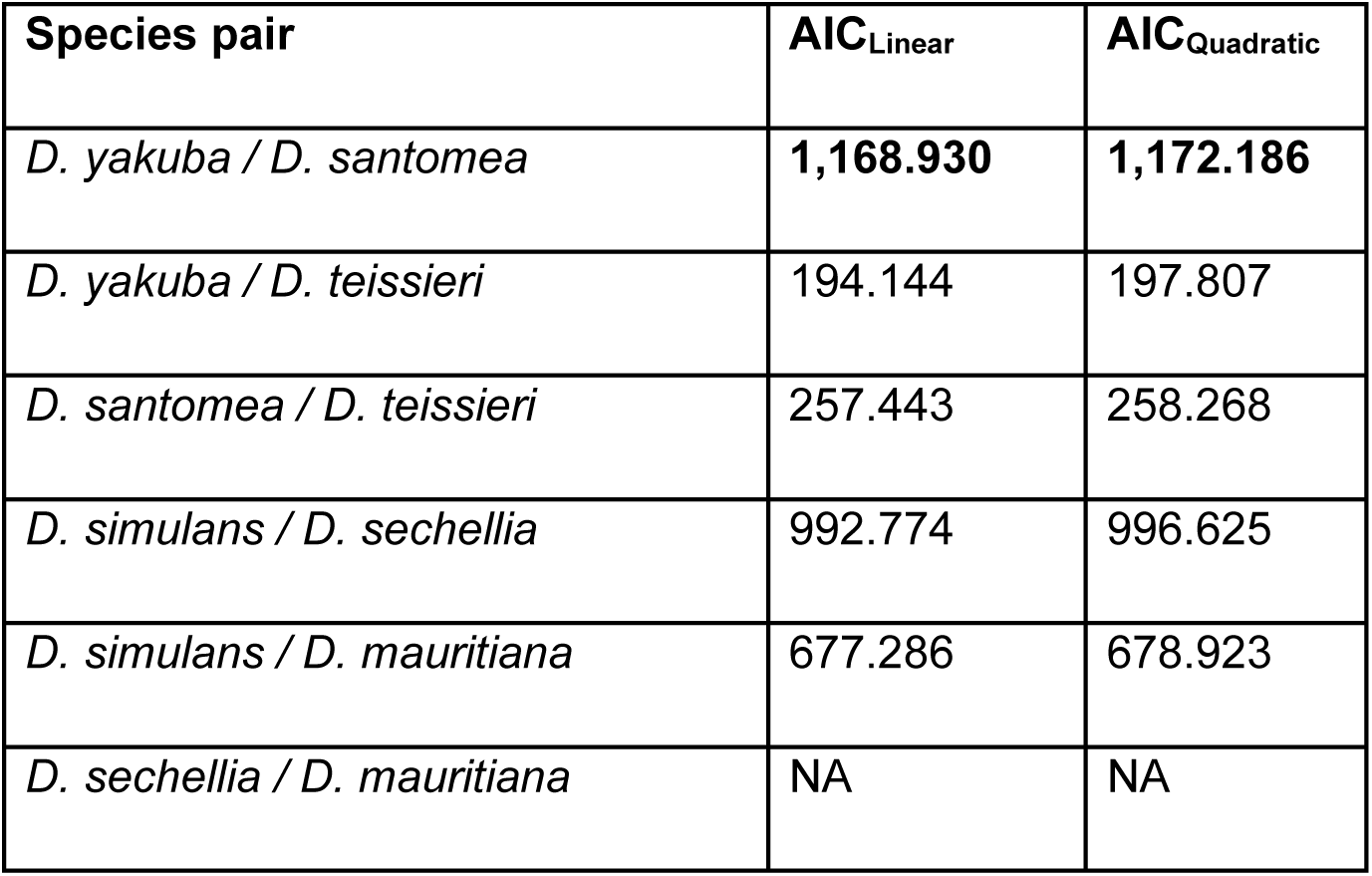
AIC values for linear and quadratic models studying the effect of temperature and reciprocal mating direction on mating latency in five heterospecifc matings.

**TABLE S12.** AIC values for linear and quadratic models studying the effect of temperature and reciprocal mating direction on mating duration in five heterospecifc matings.

| Species pair | AIC<sub>Linear</sub> | AIC<sub>Quadratic</sub> |
| --- | --- | --- |
| *D. yakuba* / *D. santomea* | 1,231.755 | 1,235.519 |
| *D. yakuba* / *D. teissieri* | 207.900 | 211.006 |
| *D. santomea* / *D. teissieri* | 253.632 | 257.381 |
| *D. simulans* / *D. sechellia* | 1,018.952 | 1,020.806 |
| *D. simulans* / *D. mauritiana* | 707.080 | 709.014 |
| *D. sechellia* / *D. mauritiana* | NA | NA |

